# A triple threat: the *Parastagonospora nodorum* SnTox267 effector exploits three distinct host genetic factors to cause disease in wheat

**DOI:** 10.1101/2021.02.25.432871

**Authors:** Jonathan K. Richards, Gayan Kariyawasam, Sudeshi Seneviratne, Nathan A. Wyatt, Steven S. Xu, Zhaohui Liu, Justin D. Faris, Timothy L. Friesen

## Abstract

*Parastagonospora nodorum* is a fungal pathogen of wheat. As a necrotrophic specialist, it deploys a suite of effector proteins that target dominant host susceptibility genes to elicit programmed cell death (PCD). Nine effector – host susceptibility gene interactions have been reported in this pathosystem, presumed to be governed by unique pathogen effectors. This study presents the characterization of the SnTox267 necrotrophic effector that hijacks two separate host pathways to cause necrosis. An association mapping approach identified *SnTox267* and the generation of gene-disrupted mutants and gain-of-function transformants confirmed its role in *Snn2*-, *Snn6*-, and *Snn7*-mediated necrosis. The *Snn2* and *Snn6* host susceptibility genes were complementary, and together they functioned cooperatively to elicit SnTox267-induced necrosis in the same light-dependent PCD pathway. Additionally, we showed that SnTox267 targeted *Snn7*, resulting in light-independent necrosis. Therefore, SnTox267 co-opts two distinct host pathways to elicit PCD. *SnTox267* sequence comparison among a natural population of 197 North American *P. nodorum* isolates revealed 20 protein isoforms conferring variable levels of virulence, indicating continuing selection pressure on this gene. Protein isoform prevalence among discrete populations indicated that SnTox267 has likely evolved in response to local selection pressures and has diversified more rapidly in the Upper Midwest. Deletion of *SnTox267* resulted in the upregulation of the unrelated effector genes *SnToxA*, *SnTox1*, and *SnTox3*, providing evidence for a complex genetic compensation mechanism. These results illustrate a novel evolutionary path by which a necrotrophic fungal pathogen uses a single proteinaceous effector to hijack two host pathways to induce cell death.

## Introduction

The plant immune system functions predominately through the recognition of microbe-produced or damage-derived molecules. This recognition of non-self leads to the induction of various defense responses. Perception of conserved microbial motifs known as pathogen-associated molecular patterns (PAMPs) can elicit a basal defense response classically known as PAMP-triggered immunity (PTI) (1). Pathogen-produced effector molecules may act to suppress or mask pathogen presence to effectively overcome PTI. However, plants have subsequently evolved receptor-like proteins to detect the presence of an effector or alternatively, the effector-mediated modification of host cellular machinery (2). Although often described as separate immune pathways, evidence suggests that PTI and ETI constituents substantially overlap and may function synergistically (3, 4). ETI operates in a gene-for-gene manner and is effective at sequestering biotrophic pathogens (2, 5, 6). Many necrotrophic pathogens, however, have evolved with their respective hosts to target the host defense response to trigger programmed cell death (PCD) and obtain nutrients from the dying tissue. This interaction has been characterized as an inverse gene-for-gene relationship where a dominant host susceptibility gene is targeted by a necrotroph-produced effector molecule (7).

*Parastagonospora nodorum* is a model necrotrophic pathogen that causes the economically important disease septoria nodorum blotch (SNB) of common wheat (*Triticum aestivum*) and durum wheat (*T. turgidum* ssp. *durum*). In contrast to the barrage of lytic enzymes often used by necrotrophic generalist pathogens, *P. nodorum* deploys an arsenal of specialized necrotrophic effector proteins that target recognition components of the host PCD pathways, resulting in necrotrophic effector triggered susceptibility. To date, nine necrotrophic effector - susceptibility gene interactions have been described, including SnToxA – *Tsn1* (8), SnTox1 – *Snn1* (9, 10), SnTox2 – *Snn2* (7), SnTox3 – *Snn3-B1*/*Snn3-D1* (11, 12), SnTox4 – *Snn4* (13), SnTox5 – *Snn5* (14), SnTox6 – *Snn6* (15), and SnTox7 – *Snn7* (16). Among these identified effector genes, *SnToxA*, *SnTox1*, and *SnTox3* have been cloned, and among the host susceptibility genes, *Tsn1*, *Snn1*, and *Snn3D-1* have been cloned (8, 10, 11, 17–19). The identified necrotrophic effector proteins exhibit typical effector properties, including the presence of a secretion signal, relatively small size (<50 kDa), and above average cysteine content. Additionally, natural populations of *P. nodorum* show varying levels of presence/absence variation of the effector loci, likely reflecting the distribution of cognate host susceptibility genes in locally planted wheat cultivars or the maintenance of beneficial alternate effector functions. These alternate functions include the ability of SnTox1 to protect from host-produced chitinases (20, 21) or SnTox3 to suppress host defense through a direct interaction with PR1 proteins (22, 23). The characterization of these effectors and corresponding host susceptibility targets has shown how *P. nodorum* specifically hijacks the host immune response and elicits cell death that ultimately provides nutrients for completion of the pathogenic life cycle.

This study shows that three previously characterized necrotrophic effectors (SnTox2, SnTox6, and SnTox7), which were originally hypothesized to function independently and target three distinct host susceptibility genes (*Snn2*, *Snn6*, and *Snn7*), are actually a single effector that targets two pathways involving three critical genes. One pathway was determined to be light-dependent and requires the cooperative function of both *Snn2* and *Snn6* to produce necrosis. The second pathway involves the host susceptibility gene *Snn7* and is postulated to be separate due to its light-independent nature. Both pathways were targeted by a novel effector, designated as SnTox267, to elicit cell death, facilitating the completion of its pathogenic life cycle. Population-specific SnTox267 protein isoforms were identified and found to vary significantly in their ability to induce necrosis. Sequence analysis also indicated that subpopulation-specific selection pressures may contribute to local *SnTox267* diversification. Additionally, the deletion of *SnTox267* resulted in the differential upregulation of effector genes *SnToxA*, *SnTox1,* and *SnTox3*, indicating the potential of intertwined effector expression networks or the existence of an underlying *trans*-acting regulatory mechanism at this genomic locus. Taken together, these results advance our understanding of complex necrotroph-host interactions.

## Results

### *Association mapping identifies a single locus associated with virulence on* Snn2 *and* Snn6

A genome-wide association study (GWAS) was conducted to identify the effectors that elicit necrosis via interaction with host susceptibility genes *Snn2* and *Snn6*. A previously described *P. nodorum* population consisting of 197 isolates from a wide geographical range in the United States (21) was used to phenotype the wheat lines BG223 (*Snn2*+) and ITMI37 (*Snn6*+) (21). We hypothesized that the use of single susceptibility gene differential lines would enable the detection of the cognate effector loci by eliminating any major background gene interactions. Phenotyping of BG223 and ITMI37 with the *P. nodorum* natural population revealed average reactions ranging from 0 to 3.5 (Fig. 1A; Supplementary Table 1). Additionally, the virulence phenotypes on the two differential lines were highly correlated (r^2^=0.59). Previously generated whole-genome sequencing reads for the natural population (NCBI BioProject PRJNA398070) were aligned to the Sn4 reference genome (24) for the identification of single nucleotide polymorphisms (SNPs) and insertions or deletions (InDels). Following filtering for minor allele frequency (5%) and missing data (30%), 322,613 polymorphic markers were identified and then used to conduct association mapping. Surprisingly, the same genomic locus on *P. nodorum* chromosome 14 was identified as significantly associated with virulence on both BG223 and ITMI37 (Fig. 1B and Fig. 1C). In both analyses, the marker SNP_66420 was the most significant association (BG223: p = 2.96 × 10^−12^; ITMI37: p = 3.23 × 10^−14^; Supplementary File 1). This marker was located 931 bp upstream of the gene *CJJ_13380*, a 798 bp single exon gene encoding a 265 amino acid protein containing 10 cysteine residues (Supplementary Figure 1). The cysteine residues were predicted to form disulfide bonds, as is commonly observed in fungal effector proteins. A signal peptide was predicted in the first 16 amino acids and the mature 27.4_kDa protein was predicted to be an effector via EffectorP (25). BLASTP searches of the NCBI non-redundant protein database failed to identify any homologous proteins, indicating that *CJJ_13380* is unique to *P. nodorum*. Two additional genes, encoding a LETM1/MDM38-like protein and a protein with unknown functional domains, were identified within the 5 kb flanking regions of SNP_66420. However, neither were predicted to encode a secreted protein and were therefore not prioritized for validation (Supplementary Figure 1).

**Figure 1.**
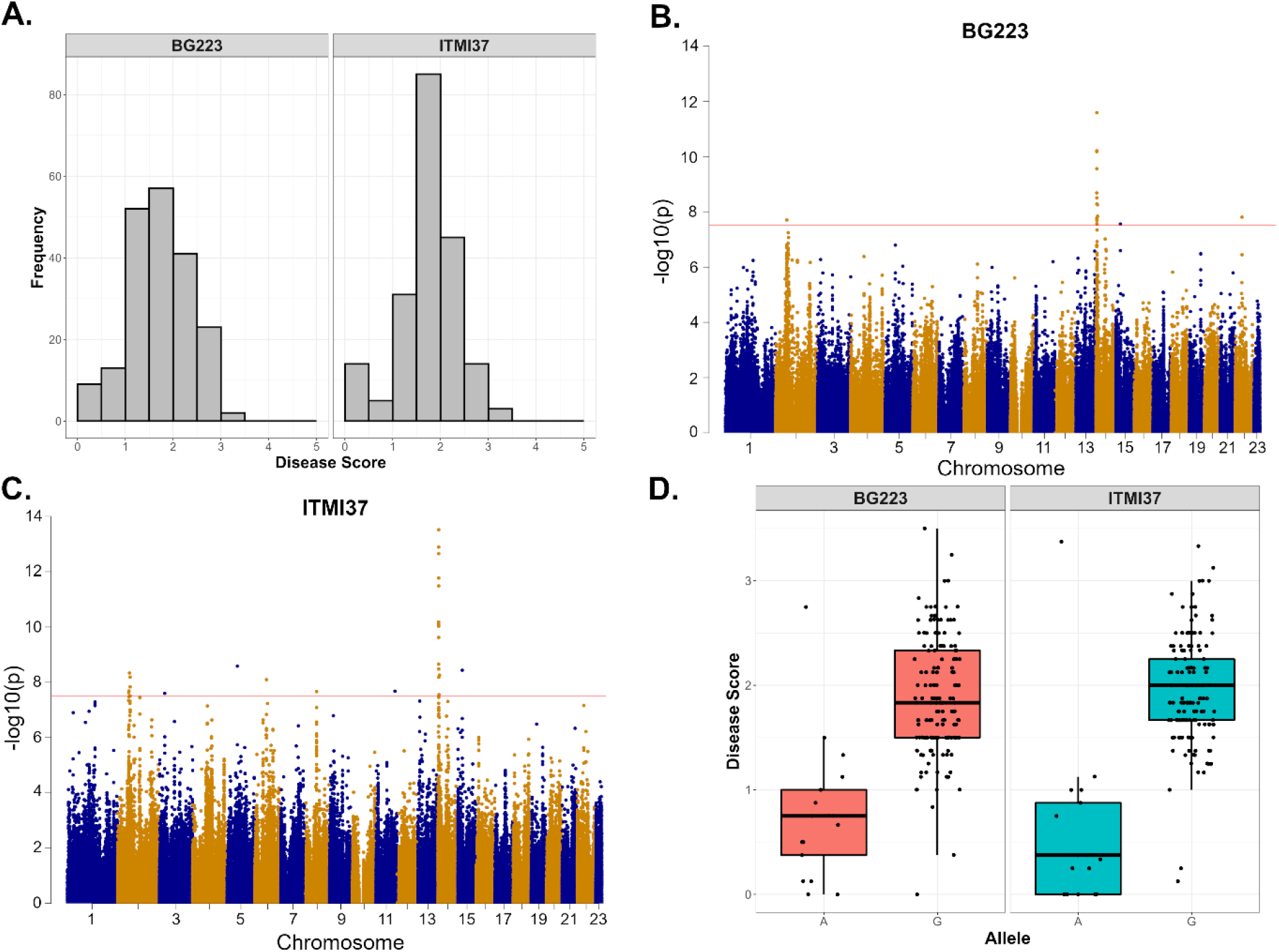
Identification of markers associated with virulence on wheat lines BG223 (*Snn2*) and ITMI37 (*Snn6*). **A**. Histogram showing the distribution of disease scores for 197 *P. nodorum* isolates on BG223 (left) and ITMI37 (right). Disease score is listed on the x-axis and frequency is listed on the y-axis. **B.** Manhattan plot illustrating markers associated with virulence on wheat line BG223. *P. nodorum* chromosomes are shown on the x-axis and marker significance is shown on the y-axis. The red line indicates a Bonferroni corrected p-value threshold of 0.01. **C.** Manhattan plot illustrating markers associated with virulence on wheat line ITMI37. *P. nodorum* chromosomes are shown on the x-axis and marker significance is shown on the y-axis. The red line indicates a Bonferroni corrected p-value threshold of 0.01. **D.** Distribution of virulence phenotypes on BG223 (left) and ITMI37 (right) for isolates that harbor the avirulent (A) and virulent (G) alleles of SNP_66420. Allele categories are listed on the x-axis and disease score is shown on the y-axis.

A total of 176 isolates had a reliable SNP_66420 genotype and two alleles were observed. Isolates having the ‘A’ allele had average disease reactions on BG223 and ITMI37 of 0.77 and 0.59, respectively (Fig. 1D). These average reaction types represent near-immune responses. Conversely, isolates harboring the ‘G’ allele exhibited average disease reactions of 1.89 and 1.96 on BG223 and ITMI37, respectively. The intermediate reaction types observed in isolates with the virulent allele could be attributed to the differential lines used harboring single susceptibility targets and the typically additive nature of necrotrophic effector – host susceptibility target interactions. These results provided the initial evidence that a single predicted effector protein was targeting at least two host susceptibility genes to elicit cell death.

### Functional validation indicates CJJ_13380 exploits three genetically distinct host factors

Following the identification of *CJJ_13380* as a strong candidate effector, we next attempted to functionally validate its role in virulence through the generation of gene-disrupted mutants. A split-marker approach was used to replace *CJJ_13380* with a hygromycin resistance gene cassette. Two independent mutants, as well as an ectopic transformant that contained a non-disrupted *CJJ_13380* and the hygromycin resistance gene inserted elsewhere in the genome, were obtained and used to inoculate the BR34 × Grandin recombinant inbred line (RIL) population (hereafter referred to as the BG population) originally used to map and characterize the *Snn2* susceptibility gene (7). Phenotyping of the *Snn2* differential line BG296 showed a marked reduction in necrosis for the Sn4ΔCJJ_13380 isolates when compared to the wild-type Sn4 and ectopic transformants (Fig. 2A). Furthermore, inoculation of the Sn4ΔCJJ_13380 deletion mutants onto the BG population, which segregates for *Snn2*, and subsequent QTL analysis failed to detect the *Snn2* QTL (Fig. 2B). Inoculation with the WT and ectopic isolates, however, identified the expected QTL corresponding to *Snn2* (Fig. 2B). To further verify the role of CJJ_13380 in *Snn2*-mediated susceptibility, the avirulent *P. nodorum* isolate Sn79-1087 was transformed with a 2086 bp genomic region containing the putative promoter and coding sequence of the candidate gene from isolate Sn4. The resulting transformed isolate was named Sn79+CJJ_13380. Inoculation of WT Sn79-1087 exhibited a near-immune response on lines containing *Snn2*, however, inoculation with the Sn79+CJJ_13380 strain showed obvious necrotic symptoms (Fig. 3A). Inoculation of the Sn79+CJJ_13380 transformant onto the BG population followed by QTL analysis identified a highly significant QTL corresponding to the *Snn2* locus (Fig. 3C). The QTL explained approximately 92% of the phenotypic variation and was the only significant locus identified. Finally, to confirm that the necrotic symptoms were due to the recognition of a secreted CJJ_13380 protein, cell free culture filtrates of Sn79+CJJ_13380 were infiltrated into leaves of the BG population and lines were evaluated for necrosis (Fig. 3B). QTL analysis again confirmed *Snn2* as the sole locus involved in this interaction (Fig. 3C). These results indicated that CJJ_13380 was a secreted protein that targeted *Snn2* to cause necrosis and was therefore SnTox2.

**Figure 2.**
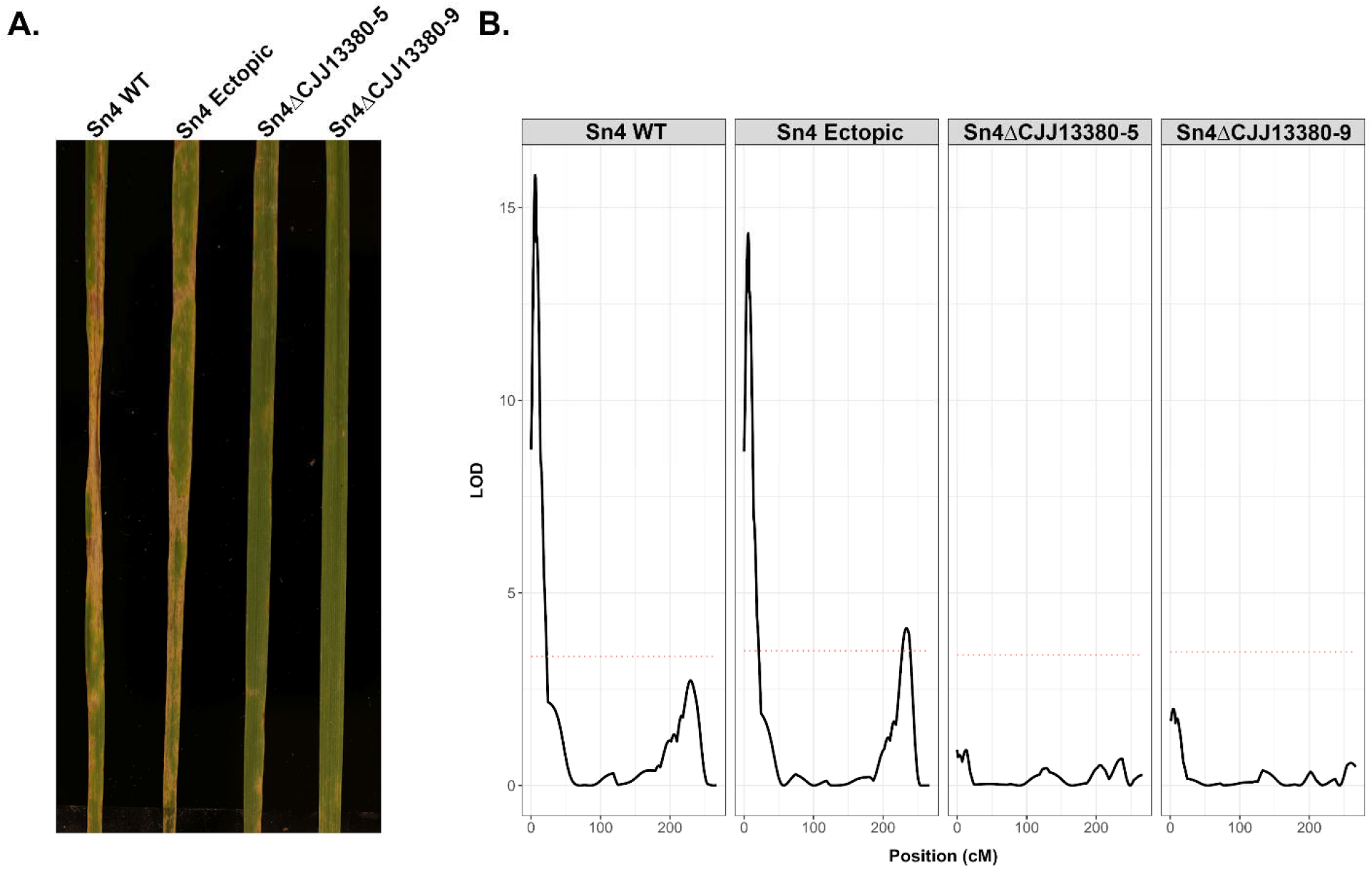
Disruption of CJJ_13380 results in the loss of virulence on lines harboring *Snn2*. **A.** Wheat line BG296 (*Snn2* differential line) inoculated with Sn4 wild-type (WT), Sn4 ectopic transformant, and two independent mutants with a disrupted *CJJ_13380* gene. **B.** QTL analysis of the BR34 × Grandin (BG) population with the Sn4 WT, Sn4 ectopic, and two CJJ_13380-disrupted isolates. Genetic positions along wheat chromosome 2D are listed in cM on the x-axes and the logarithm of the odds (LOD) is shown on the y-axis. The red dotted line corresponds to a LOD threshold of p=0.05 as determined by 1000 permutations conducted for each analysis separately.

**Figure 3.**
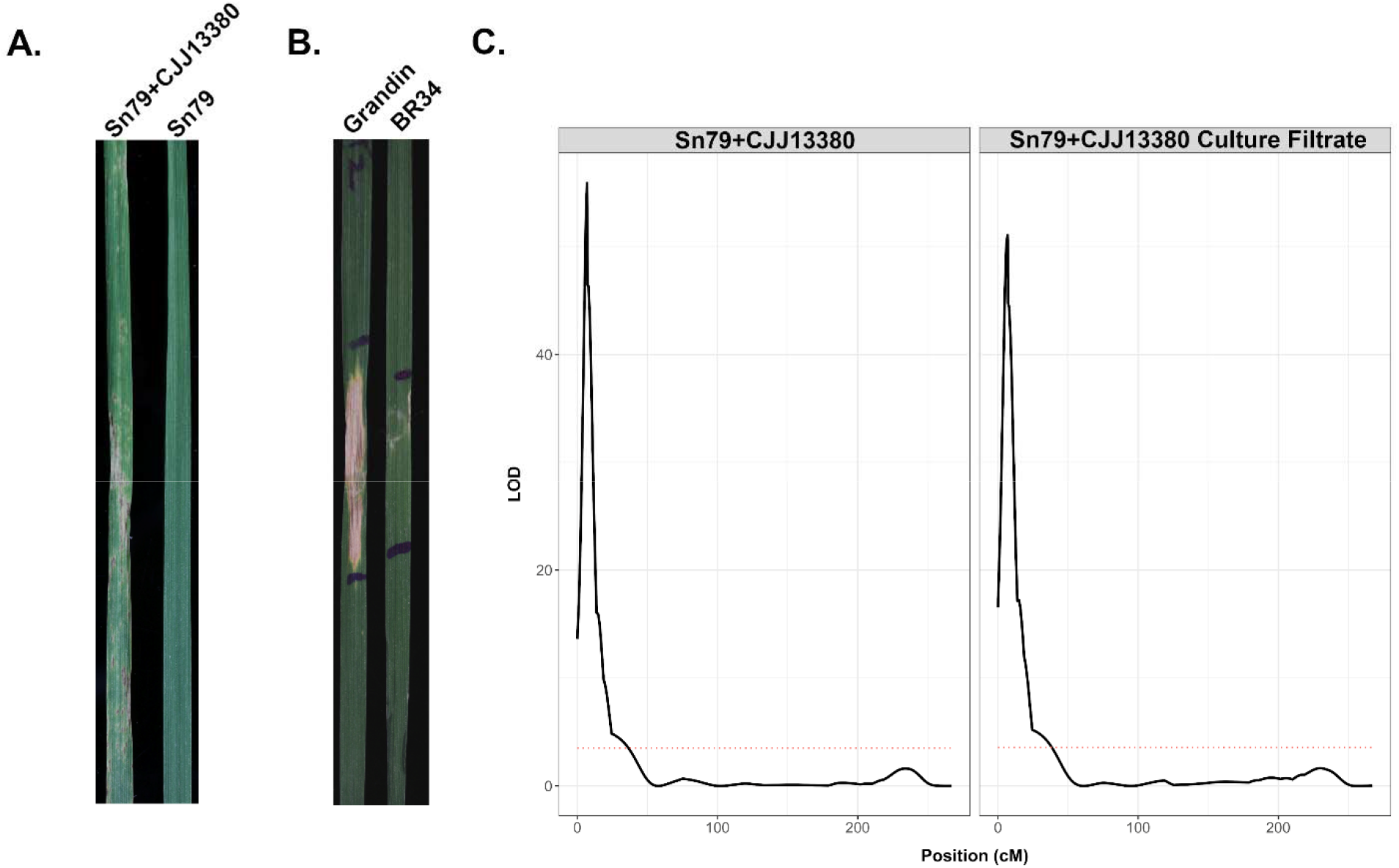
Inoculation of gain-of-function transformant Sn79+CJJ13380 and infiltration of culture filtrates result in virulence or cell death on lines harboring *Snn2*. **A.** Inoculation of BG296 (*Snn2* differential line) with gain-of-function transformant Sn79+CJJ13380 and Sn79 WT. **B.** Infiltration of Sn79+CJJ13380 culture filtrates on wheat lines Grandin (sensitive) and BR34 (insensitive). **C.** QTL analysis of the BR34 × Grandin (BG) population with the Sn79+CJJ13380 isolate (left) and infiltration of Sn79+CJJ13380 culture filtrates (right). Genetic positions along wheat chromosome 2D are listed in cM on the x-axes and the logarithm of the odds (LOD) is shown on the y-axis. The red dotted line corresponds to a LOD threshold of p=0.05 as determined by 1000 permutations conducted for each analysis separately.

Because the identical SNP in the putative promoter of *CJJ_13380* was also associated with virulence on ITMI37, the *Snn6* differential line, we used the same deletion and expression transformants to inoculate the International Triticeae Mapping Initiative (ITMI) population, which segregated for *Snn6*. Inoculation of the Sn4ΔCJJ_13380 knockout mutants onto the *Snn6* differential line ITMI37 showed a marked reduction in necrotic symptoms compared to inoculations with Sn4 WT and the Sn4 ectopic transformant (Fig. 4A). Inoculation of the Sn4 WT and Sn4 ectopic transformant onto the ITMI population detected a QTL corresponding to *Snn6*. However, inoculation of Sn4ΔCJJ_13380 mutants onto the ITMI population and subsequent QTL analysis failed to detect significance at the *Snn6* locus (Fig. 4B). Additionally, inoculation of the Sn79+CJJ_13380 isolate onto ITMI37 showed typical disease development, whereas ITMI37 inoculations with Sn79-1087 WT exhibited an immune response (Fig. 5A). Moreover, inoculation of Sn79+CJJ_13380 onto the ITMI wheat population detected a single, highly significant QTL corresponding to *Snn6* that explained approximately 42.13% of the phenotypic variation (Fig. 5B). Finally, infiltration of culture filtrates of the Sn79+CJJ_13380 isolate into leaves of the ITMI population also mapped sensitivity to the *Snn6* locus and explained 54.67% of the variation (Fig. 5C). These results indicate that in addition to eliciting necrosis via *Snn2*, CJJ_13380 also targets *Snn6* to cause necrosis.

**Figure 4.**
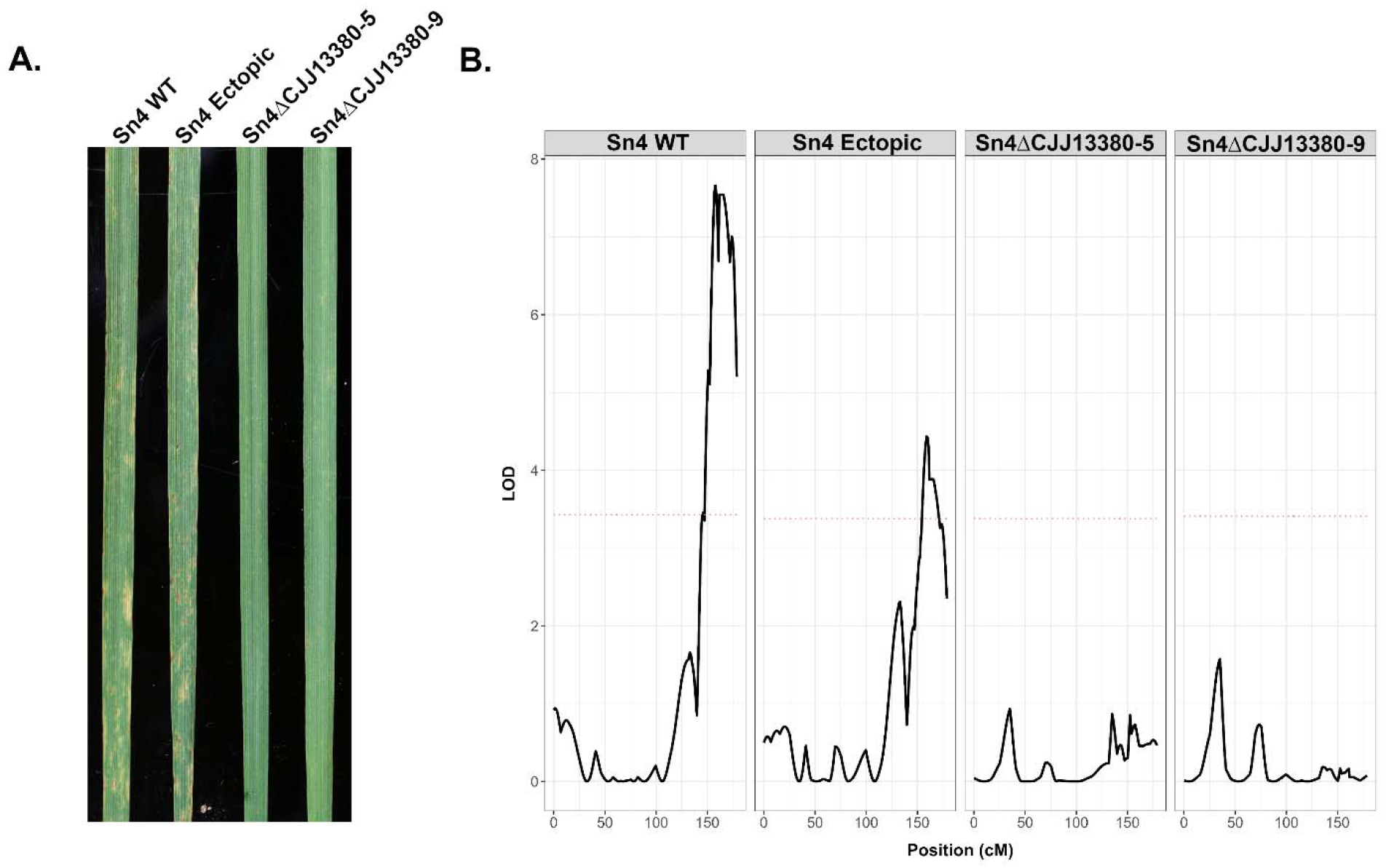
Disruption of CJJ_13380 results in the loss of virulence on lines harboring *Snn6*. **A.** Wheat line ITMI37 (*Snn6* differential line) inoculated with Sn4 wild-type (WT), Sn4 ectopic transformant, and two independent mutants with a disrupted *CJJ_13380* gene. **B.** QTL analysis of the ITMI population with the Sn4 WT, Sn4 ectopic, and two CJJ_13380-disrupted isolates. Genetic positions along wheat chromosome 6A are listed in cM on the x-axes and the logarithm of the odds (LOD) is shown on the y-axis. The red dotted line corresponds to a LOD threshold of p=0.05 as determined by 1000 permutations conducted for each analysis separately.

**Figure 5.**
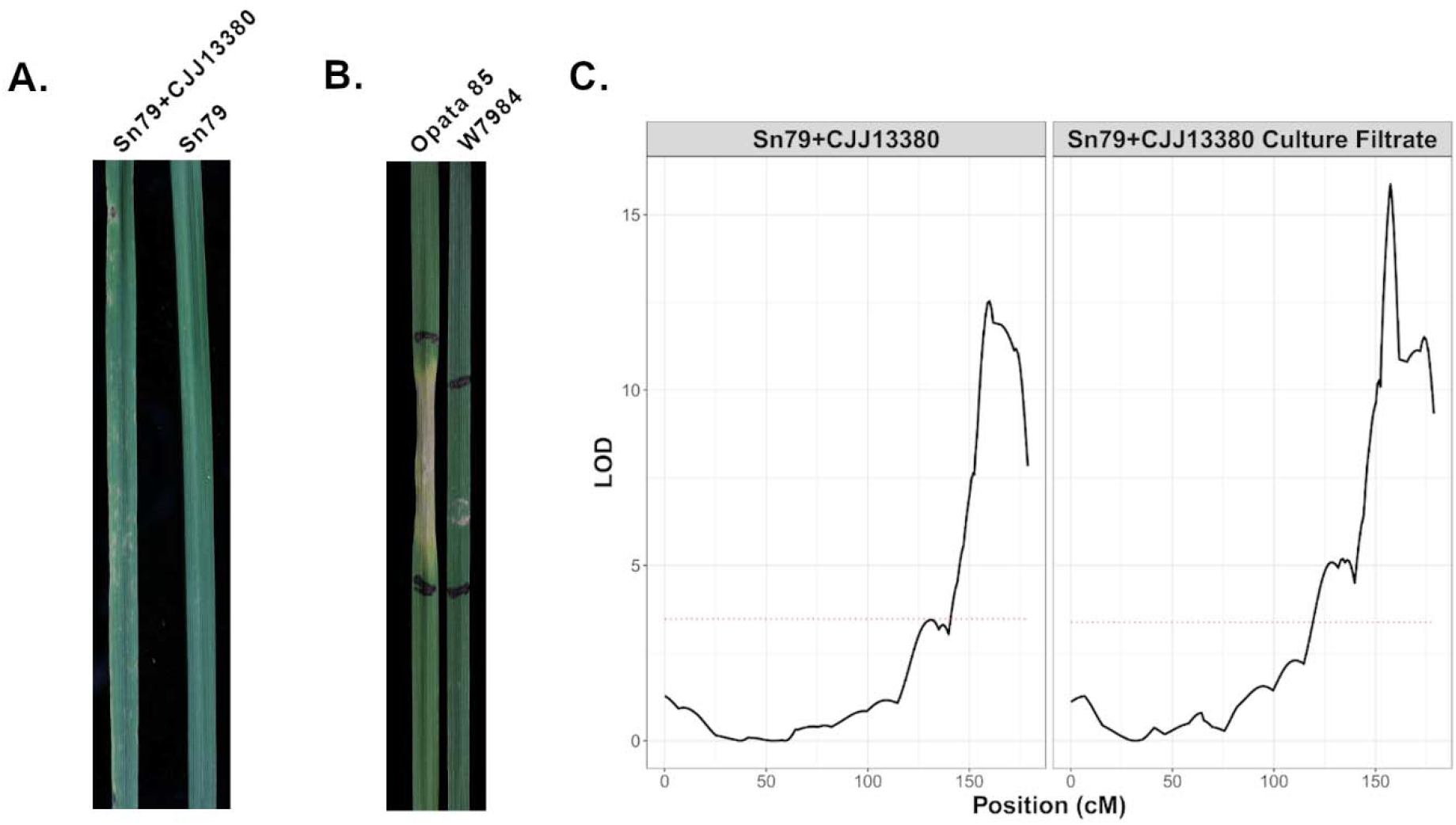
Inoculation of gain-of-function transformant Sn79+CJJ13380 and infiltration of culture filtrates result in virulence or cell death on lines harboring *Snn6*. **A.** Inoculation of ITMI37 (*Snn6* differential line) with gain-of-function transformant Sn79+CJJ13380 and Sn79 WT. **B.** Infiltration of Sn79+CJJ13380 culture filtrates on wheat lines Opata 85 (sensitive) and W7984 (insensitive). **C.** QTL analysis of the ITMI population with the Sn79+CJJ13380 isolate (left) and infiltration of Sn79+CJJ13380 culture filtrates (right). Genetic positions along wheat chromosome 6A are listed in cM on the x-axes and the logarithm of the odds (LOD) is shown on the y-axis. The red dotted line corresponds to a LOD threshold of p=0.05 as determined by 1000 permutations conducted for each analysis separately.

Previous research mapped the dominant susceptibility gene *Snn7*, which was targeted by the SnTox7 effector, to chromosome 2D in a Chinese Spring (CS) × CS-Timstein 2D (substitution of CS chromosome 2D by Timstein chromosome 2D) population (16). The characteristics derived from previous partial purification of SnTox7 including an estimated size of 10 – 30 kDa, were similar to those observed for the partially purified SnTox2 and SnTox6 (7, 15), as well as the effector identified in this study. Therefore, we tested the potential for this novel effector to elicit cell death via targeting of *Snn7*. A CS × CS-Tm 2D F_2_ population was developed to verify whether the *CJJ_13380* candidate gene was also targeting *Snn7* on chromosome 2D. Nine markers specific to chromosome 2D were used for genotyping and development of the corresponding linkage group. The F_2_ population was infiltrated with culture filtrates of Sn79+CJJ_13380 and QTL analysis revealed a highly significant peak corresponding to the *Snn7* locus, indicating that *Snn7* is also targeted by CJJ_13380 and conditions sensitivity to it (Supplementary Fig. 2).

Taken together, these results indicated that CJJ_13380 was a single secreted necrotrophic effector protein that elicited cell death from wheat lines harboring *Snn2*, *Snn6*, or *Snn7*. We propose the effector nomenclature *SnTox267* to reflect the three known host sensitivity/susceptibility targets.

### Snn2 *and* Snn6 *function cooperatively to mediate SnTox267 elicited necrosis*

Because SnTox267 induced necrosis on lines harboring *Snn2*, *Snn6*, or *Snn7*, we wanted to determine if the host targets represented distinct pathways or were polymorphic components of a single pathway. Previously, the *Snn2* – *SnTox2* and *Snn6 – SnTox6* interactions were shown to be light dependent, whereas the *Snn7* – *SnTox7* interaction was shown to be light independent and therefore likely functions in a separate pathway (7, 15, 16). We also confirmed in this study, the light dependency or independency of these reactions using side-by-side SnTox267 infiltrations (Supplementary Fig. 3). Therefore, we specifically examined the potential of *Snn2* and *Snn6* functioning in the same pathway. An F_2_ population (n=159) was developed from a cross between the SnTox267-sensitive lines BG223 (*Snn2* differential line) and Opata 85 (sensitive parent of the ITMI population and possesses *Snn6*) to test whether *Snn2* and *Snn6* are complementary genes. If *Snn2* and *Snn6* elicited cell death independently, we expected an F_2_ segregation ratio of 15:1 (sensitive:insensitive) for two dominant genes with redundant functions. However, all 159 F_2_ progeny were sensitive to SnTox267 infiltrations. This indicated that BG223 and Opata85 each harbored functional copies of both *Snn2* and *Snn6*, therefore, all F_2_ progeny were fixed at both loci. Based on these results, we hypothesized that both *Snn2* and *Snn6* are required for SnTox267 sensitivity. If this model was correct, the insensitive parents of the BG and ITMI populations would harbor a functional copy of one susceptibility gene (either *Snn2* or *Snn6*) and a non-functional copy of the remaining susceptibility gene, which compromises the pathway and leads to insensitivity. Crossing the two insensitive (resistant) lines would then restore the functional pathway in the progeny and result in SnTox267 sensitivity. To test this hypothesis, we phenotyped F_1_ and F_2_ progeny of a cross between BR34 (insensitive parent of the BG population) and W7984 (insensitive parent of the ITMI population) by infiltrating Sn79+SnTox267 culture filtrate. All F_1_ progeny were sensitive (n=3; Fig. 6) and sensitivity in the F_2_ individuals (n=49) segregated 29:20 (sensitive:insensitive), fitting the expected 9:7 ratio for two genes with complementary gene action (χ^2^ goodness-of-fit test: χ^2^=0.17, p=0.68; Fig. 6). Taken together, these results indicated that *Snn2* and *Snn6* functioned as two components of the same molecular pathway where both dominant genes were required for necrosis development induced by SnTox267.

**Figure 6.**
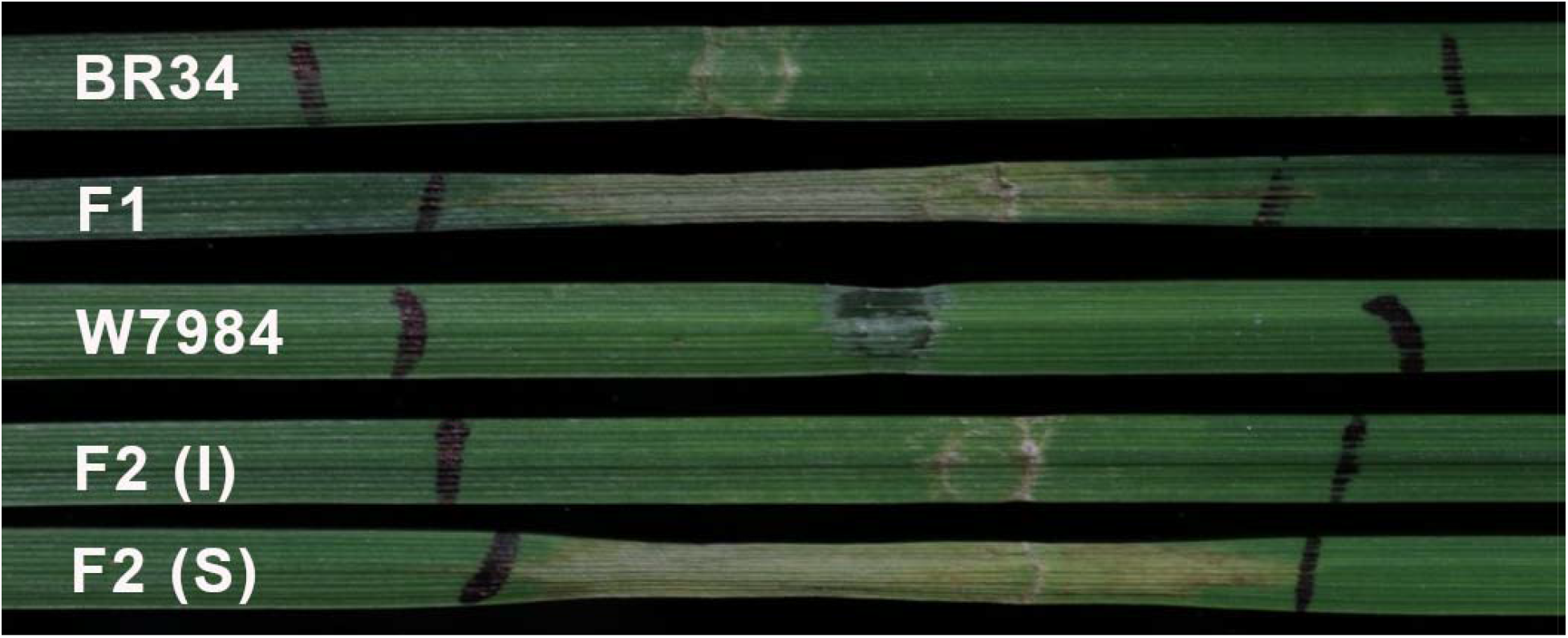
*Snn2* and *Snn6* are both required to produce SnTox267-mediated cell death. Culture filtrates of the gain-of-function transformant Sn79+CJJ13380 were infiltrated in wheat lines BR34 (*snn2*/*Snn6*), F_1_ plants (n=3) derived from a cross of BR34 and W7984, W7984 (*Snn2*/*snn6*), and F_2_ (n=49) plants derived from a cross of BR34 and W7984. ‘I’ indicates insensitive and ‘S’ indicates sensitive reactions.

### SnTox267 exhibits an uneven distribution of nucleotide diversity

Fungal effector genes, including the previously cloned *SnToxA, SnTox1,* and *SnTox3*, have the propensity to exhibit presence/absence variation. BLAST searches of *de novo* assembled genomes from each isolate in the natural population (n=197) revealed that a total of 95.43% (188/197) of isolates had full or partial *SnTox267* sequence represented in their respective *de novo* assemblies (deposited at doi:10.5281/zenodo.4560540). These results indicated that *SnTox267* is only absent from a small proportion of the natural population.

Sequence variants in the *SnTox267* coding region were extracted and resulted in the identification of 22 polymorphic nucleotides, corresponding to an overall nucleotide diversity of 0.005. A higher level of nucleotide diversity was observed among isolates from the Upper Midwest population compared to the Southern/Eastern population with population-specific nucleotide diversity values of 0.006 and 0.002, respectively. Among the identified SNPs, six were synonymous changes, 15 were non-synonymous changes, and one introduced a premature stop codon (Fig. 7A). Sliding window analysis of nucleotide diversity determined that variation was not uniformly distributed and was generally concentrated in the 3’ half of the gene (Fig. 7B). Taken together, these results indicated that the presence of *SnTox267* was largely conserved within a diverse *P. nodorum* natural population, however, it possessed considerable nucleotide diversity that was localized to the 3’ portion of the gene.

**Figure 7.**
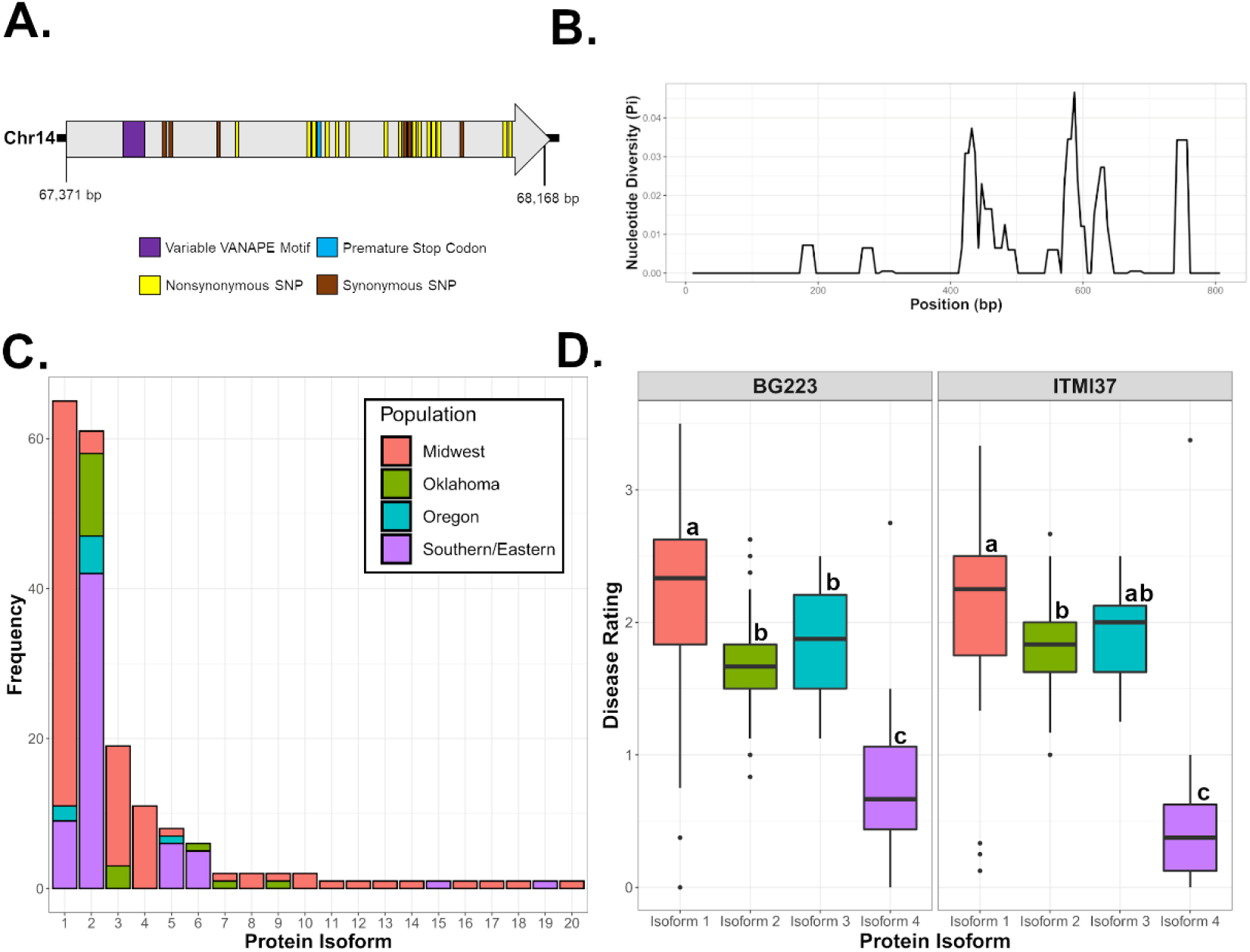
Sequence diversity of *SnTox267* in a natural population (n=197) of *P. nodorum*. **A.** Gene model of *SnTox267* located on chromosome 14. The arrow corresponds to the single exon structure of *SnTox267*. Colored boxes illustrate the locations of polymorphisms (annotated in legend). **B.** Nucleotide diversity within the *SnTox267* coding region. The position (bp) is shown on the x-axis and the nucleotide diversity (π) is shown on the y-axis. **C.** Prevalence of protein isoforms in discrete *P. nodorum* populations. The specific isoforms are listed on the x-axis and the frequency of each isoform is shown on the y-axis. Colors shown in the legend (upper right) correspond to discrete populations previously determined by (21). **D.** Quantitative virulence profiles of protein isoform groups with significant representation (n>10). The specific isoforms are listed on the x-axis and disease ratings are shown on the y-axis. Letter codes signify significantly different groups as determined by a Dunn’s multiple comparisons test (Bonferroni adjusted p-value < 0.10).

### SnTox267 protein isoform diversity varies between populations and contributes to virulence

Within the entire population, a total of 32 nucleotide haplotypes, considering SNPs and InDels, were identified in SnTox267 with a haplotype diversity (H_d_) of 0.86. This level of haplotype diversity indicated a high probability that two randomly chosen isolates would possess different haplotypes. However, a large proportion of the population (30.32%, n=57) comprised a single nucleotide haplotype. Examination of translated protein sequences of each haplotype resulted in a total of 20 protein isoforms with two predominate isoforms comprising 34.57% (Isoform 1) and 32.45% (Isoform 2) of the population. Only two amino acid substitutions differentiated these two isoforms, an aspartic acid to alanine substitution at residue 137 (D137A) and a threonine to serine substitution at residue 244 (T244S; Supplementary Fig. 4). Interestingly, the prevalence of these two isoforms varied substantially between the previously defined *P. nodorum* populations (21). Among isolates harboring the SnTox267 Isoform 1, 83.08% were from the Upper Midwest (54.5% of all Upper Midwest isolates), 13.85% were from the Southern/Eastern United States (14.1% of all Southern/Eastern isolates), and 3.08% from Oregon (25.0% of all Oregon isolates). In contrast, among isolates with SnTox267 Isoform 2, 68.85% of isolates were from the Southern/Eastern United States (65.6% of all Southern/Eastern isolates), 18.03% from Oklahoma (68.8% of all Oklahoma isolates), 8.20% from Oregon (62.5% of all Oregon isolates), and only 4.92% from the Upper Midwest (3.0% of all Upper Midwest isolates). Large differences were also observed in the total number of isoforms present in discrete populations. The Upper Midwest population contained 17 protein isoforms, whereas the Southern/Eastern population only had six isoforms present (Fig. 7C). This stark contrast in the total number of isoforms detected in each population suggests SnTox267 has been evolving more rapidly or over a longer evolutionary period in the Upper Midwest compared to the Southern/Eastern United States. The differences in geographical prevalence of the predominate isoforms also indicated that local selection pressures were actively driving the maintenance of specific isoforms in each population, potentially due to allelic differences in cognate host susceptibility genes.

Virulence phenotypes of isolates harboring specific protein isoforms were examined to determine the potential effects of non-synonymous substitutions. A protein isoform was considered to confer virulence if disease reactions on BG223 and ITMI37 were greater than 1.0 and only isoforms with sufficient representation were compared (n>10). Three of the isoforms were considered virulent, with average disease reactions on BG223 and ITMI37 ranging from 1.66 – 2.17 and 1.80 – 2.11, respectively (Fig. 8D). Significant differences in virulence between these isoforms were detected, indicating that the subtle mutations differentiating them may have quantitatively contributed to virulence (Dunn’s multiple comparison test, Bonferroni adjusted p-value < 0.10). A premature stop codon was identified in isoform 4 resulting in average disease reactions on BG223 and ITMI37 of 0.85 and 0.63, respectively (Fig. 7D). Although the remaining 16 isoforms are represented by fewer than 10 isolates per isoform and were therefore not included in statistical comparisons, it is worth noting that 11 isoforms appeared to confer virulence, with average disease reactions on BG223 and ITMI37 ranging from 1.33 – 2.42 and 1.44 – 2.17, respectively (Supplementary Table 2). The remaining five isoforms, all represented by single isolates from the Upper Midwest, were considered avirulent with average disease reactions on BG223 and ITMI37 ranging from 0 – 0.875 and 0 – 1.125, respectively. Two of these isoforms contained premature stop codons indicating a truncated protein likely led to the loss of virulence. Two isoforms harbored a six amino acid deletion (discussed further below) that likely disrupted protein function. The remaining avirulent isoform was found to have two non-synonymous mutations that differentiated it from the rest of the isoforms. One mutation induced a phenylalanine to tyrosine substitution at residue 187 (F187Y) and the other caused a threonine to alanine substitution at residue 244 (T244A). Interestingly, valine and serine substitutions were also identified in three and six isoforms, respectively, at residue 244. The T244S substitution had no apparent effect on virulence, as these isoforms were all classified as virulent. Three isoforms had the T244V substitution and were avirulent, however, two of these isoforms had premature stop codons. The changes in amino acids at this position may have altered the phenotype due to changes in hydrophobicity. As threonine is a hydrophilic amino acid, a substitution to serine, also a hydrophilic amino acid, may have minimal effect. The substitutions to hydrophobic residues such as alanine and valine may have led to altered effector activity. Although represented by a small sample size, these results indicated that the F187Y, T244A, or T244V mutations may have led to a loss of virulence, but further experiments are warranted to independently validate these effects.

**Figure 8.**
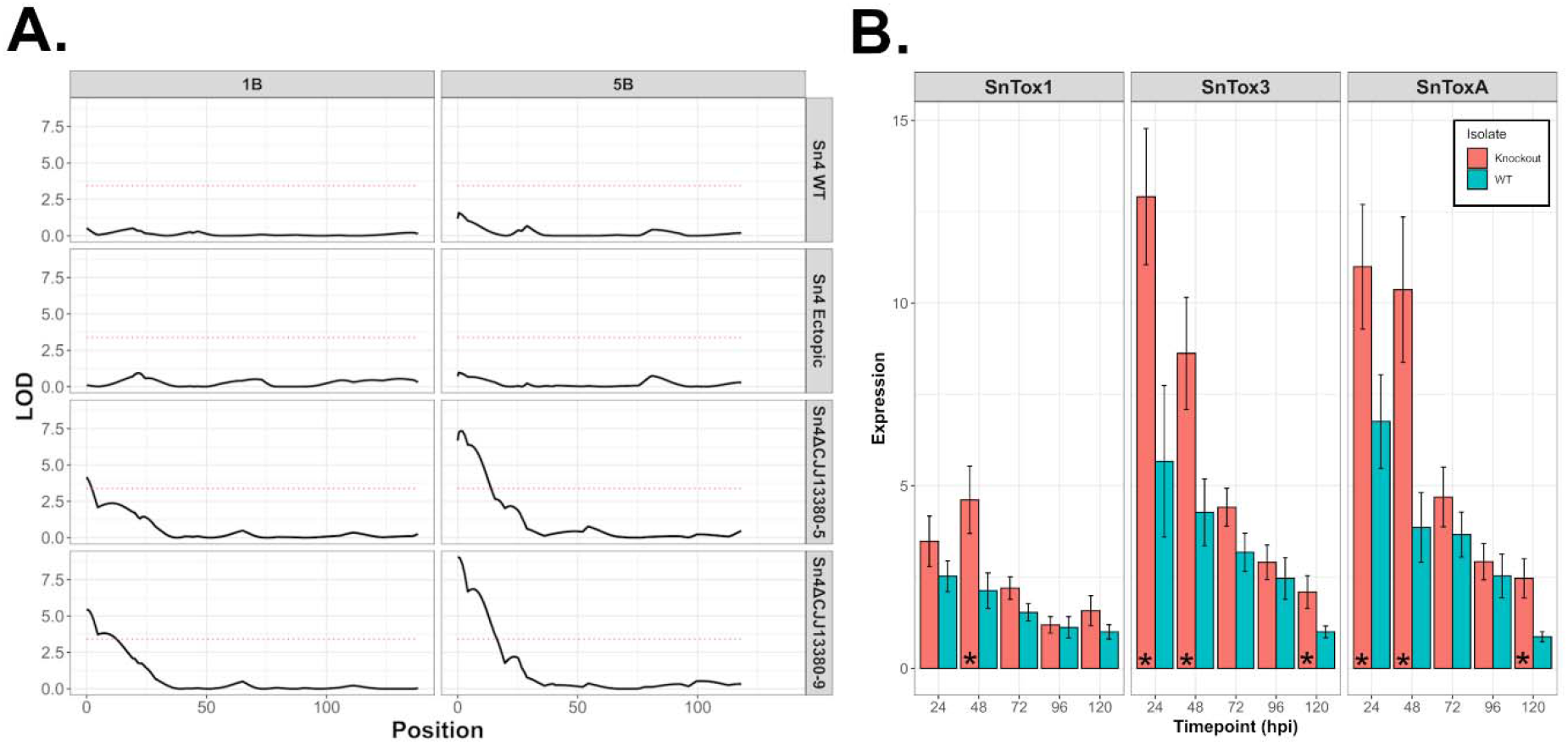
Disruption of *SnTox267* resulted in the upregulation of unrelated effectors, leading to detection of corresponding susceptibility loci. **A.** QTL analysis of the ITMI population inoculated with Sn4 WT, Sn4 ectopic transformant, and two Sn4_Δ_CJJ1380 mutants. QTL corresponding to *Snn1* on wheat chromosome 1B (left) and *Snn3* on chromosome 5B (right) were only detected following inoculation with the gene-disrupted mutants. Genetic positions (cM) are listed on the x-axes and the logarithm of the odds (LOD) values are shown on the y-axes. The red dotted line corresponds to a LOD threshold of p=0.05 as determined by 1000 permutations conducted for each analysis separately. **B.** A qPCR assay to evaluate differences in *SnTox1*, *SnTox3*, and *SnToxA* expression at five *in planta* timepoints (24, 48, 72, 96, and 120 hours post-inoculation). Timepoints are shown on the x-axes and relative expression values are shown on the y-axis. Bar colors correspond to either the WT or knockout strain Sn4_Δ_CJJ13380-5 (legend on the right). Error bars represent the standard error of the means (SEM). ‘*’ represents significant upregulation in the knockout compared to the WT at the same timepoint (Student’s t-test, p<0.05).

Alignments of the protein isoform amino acid sequences revealed a variable amino acid repeat stemming from an 18 bp in-frame insertion or deletion (Supplementary Fig. 4; Supplementary Table 2). The 18 bp insertion/deletion translated to a VANAPE motif that was predominantly present in two tandem copies in 91.4% of isolates. Three isolates possessed isoforms with three tandem copies, while isoforms from 14 isolates only harbored one copy. Among the 14 isolates harboring isoforms with only one copy of the VANAPE motif, 12 also belonged to haplotypes containing a premature stop codon. The two remaining isolates that possessed the single VANAPE motif without a nonsense mutation had markedly lower virulence of 0.375 on both BG223 and ITMI37. However, due to a small sample size, it is difficult to say with any certainty that this deletion alone leads to compromised effector function.

### *Disruption of* SnTox267 *results in compensatory upregulation of* SnToxA, SnTox1, *and* SnTox3

Following confirmation that *SnTox267* functioned in the elicitation of necrosis from lines containing *Snn2*, *Snn6*, and *Snn7*, we wanted to understand the transcriptional regulation during infection timepoints. An RT-qPCR experiment was conducted using the Sn4 WT isolate inoculated onto the *Snn2* line Grandin. Timepoints assayed included 24, 48, 72, 96, and 120 hours post-inoculation (hpi). *SnTox267* expression was normalized against the housekeeping actin gene. Expression appeared to peak at 24 hpi and slowly declined throughout the timepoints examined, indicating that *SnTox267* is activated and functions in early infection (Supplementary Fig. 5).

*P. nodorum* isolate Sn4 harbors functional copies of several characterized NEs in addition to *SnTox267*, including *SnTox1*, *SnTox3*, and *SnToxA*. However, QTL analysis following the inoculation of Sn4 WT and the Sn4 ectopic transformant onto the ITMI population, which also segregates for *Snn1* and *Snn3*, did not detect either *Snn1* or *Snn3* as significantly associated with disease (Fig. 8A). Interestingly, inoculations with both Sn4ΔSnTox267 mutants resulted in the detection of *Snn1* and *Snn3* (Fig. 9A). We hypothesized that the deletion of *SnTox267* may alter the expression levels of additional effector genes, including *SnToxA, SnTox1*, and *SnTox3*, as a compensatory measure to account for the loss of an important effector.

To test the hypothesis that expression of *SnTox1, SnTox3,* and *SnToxA* is altered in the Sn4ΔSnTox267 isolate, expression patterns of these effector genes were compared to the Sn4 WT isolate using qPCR at 24, 48, 72, 96, and 120 hpi. Expression of each gene was measured relative to the fungal actin gene. Our results showed that expression of all three genes peaked at 24 hpi in Sn4 WT. Similarly, *SnTox3* and *SnToxA* expression in Sn4ΔSnTox267 peaked at 24 hpi, however, expression of *SnTox1* peaked at 48 hpi.

When comparing expression of each effector gene between Sn4 WT and Sn4ΔSnTox267, higher expression of *SnToxA*, *SnTox1*, and *SnTox3* was observed in Sn4ΔSnTox267 at each timepoint compared to the Sn4 WT (Fig. 8B). The expression of the three effectors in Sn4ΔSnTox267 was significantly higher compared to that of Sn4 WT during the first 48h of the infection process. Specifically, expression of *SnToxA* and *SnTox3* was significantly upregulated at 24 and 48 hpi, while *SnTox1* was significantly upregulated at 48 hpi. Additionally, expression of *SnToxA* and *SnTox3* appeared to be sustained longer, as these genes were significantly upregulated at 120 hpi (Figure 8B). These results indicated that *P. nodorum* was attempting to compensate for the loss of *SnTox267* in Sn4ΔSnTox267 by the upregulation of other genetically unlinked necrotrophic effector genes during the initial stages of the infection process.

## Discussion

In this study, we identified the effector SnTox267 which exploits two distinct host pathways involved in the elicitation of cell death and disease. This represents a novel evolutionary route whereby a necrotrophic pathogen uses a single protein to hijack three genetically distinct pieces of host cellular machinery from two apparently separate physiological pathways to induce PCD. Our results also indicate that the two host susceptibility genes, *Snn2* and *Snn6*, are acting cooperatively in the same pathway to mediate necrosis elicited by a single effector. Currently, the host targets are not characterized, although this study lays the foundation for the dissection of the distinct pathways leading to SnTox267-mediated necrosis.

The functional characterization of SnTox267 and genetic dissection of host susceptibility has further clarified our understanding of inverse gene-for-gene interactions. Previously, the *Snn2*, *Snn6*, and *Snn7* genes were thought to function as susceptibility targets via independent interactions with unique effectors (7, 15, 16). However, the results from this study indicate that a single effector has co-opted these host targets or pathways to elicit cell death. From a fundamental perspective, this novel adaptive mechanism illustrates the plasticity of effector-producing fungal pathogens and their ability to use multiple evolutionary scenarios to their advantage. A similar, yet distinctly different interaction has been identified in the *P. nodorum* – wheat pathosystem with the homoeologous loci *Snn3-B1* and *Snn3-D1*. Previously, the *P. nodorum* effector SnTox3 was observed to interact with two host sensitivity genes, *Snn3-B1* and *Snn3-D1*, which were homoeologous and located on chromosome 5B of the B-genome containing polyploids and 5D of the diploid D-genome progenitor *Aegilops tauschii* (12). *Snn3-B1* and *Snn3-D1* were derived from a common ancestor and evolved independently without selection pressure before the introduction of *SnTox3* (12). Therefore, both loci maintained effector recognition capabilities. Although representative of multiple host targets, the SnTox3-Snn3-B1/D1 interactions are inherently different than the reported SnTox267 interactions. The homoeologous locations of the *Snn3* genes and phenotypic similarities of the induced necrosis point to a single host pathway being exploited. In contrast, the non-homoeologous positions of the SnTox267 susceptibility loci, as well as the variable light dependency of SnTox267-induced necrosis, illustrate a different scenario where the pathogen has taken multiple evolutionary steps to hijack host cell death pathways.

Effector recognition by two host genes resulting in cell death has been previously reported in the hemibiotrophic *Leptosphaeria maculans* – oilseed rape pathosystem. The effector AvrLm4-7 is recognized by both the *Rlm4* and *Rlm7* resistance genes (26) where a single amino acid change governs recognition specificity by *Rlm4* while *Rlm7* retains the ability to recognize either isoform. Additionally, the *L. maculans* effector AvrLm1 is recognized by both host resistance genes *Rlm1* and *LepR3*, the latter being cloned and characterized as a receptor-like kinase (27, 28). In contrast to the inverse gene-for-gene relationship of *SnTox267* with its cognate susceptibility genes, these unique examples of gene-for-gene interactions result in an evolutionary advantage for the host, because the recognition of the pathogen produced effectors results in a defense response that impedes the pathogen.

As the host targets of SnTox267 are presently unidentified, it was initially unclear if *Snn2*, *Snn6*, and *Snn7* represented translocated paralogs, nonhomologous genes with common structural motifs targeted by SnTox267, or genes within the same pathway. The observed complementary gene action of *Snn2* and *Snn6* support our hypothesis that these two genes function cooperatively in the same pathway. It is expected that host gene diversification leading to escape of effector targeting in a necrotrophic system (i.e. resistance) would occur in genes involved in direct or indirect effector interactions rather than downstream signaling components which typically remain conserved due to potential roles in other physiological processes (29). Therefore, the possibility exists that both *Snn2* and *Snn6* are involved in effector interaction, with SnTox267 potentially hijacking a two-component system operating in a guard or decoy model (30). In these scenarios, a nucleotide-binding leucine-rich repeat receptor (NLR) guards the effector target or a decoy target. Once modification of, or recognition by the target is sensed by the NLR, a resistance response is triggered, resulting in PCD. More recently, the requirement of two host NLR genes, one of which possesses an integrated domain, to confer resistance has been observed in several pathosystems (31–35). In these cases, the resistance genes are genetically colocalized and disruption of either member of the resistance gene pairs compromises resistance. The integrated domains of NLRs are widespread in plant species and are postulated to act as decoy targets or sensors, but also may retain biochemical or functional activity (36–38). Therefore, a possible hypothesis is that *Snn2* and *Snn6* are two functional susceptibility genes working cooperatively in a guard or decoy model, analogous to the aforementioned resistance genes, with the key difference being the triggering of necrotrophic effector triggered susceptibility rather than effector triggered immunity.

The light independent nature of *Snn7*-mediated necrosis points to the existence of a second pathway exploited by SnTox267. Although the precise mechanisms that govern light dependency in effector-mediated cell death reactions are not fully understood, previous research has implicated light-dependency as requisite for specific effector-induced cell death responses (39, 40). In the *P. nodorum* – wheat pathosystem, the previously characterized effectors SnToxA, SnTox1, and SnTox3 were demonstrated to induce necrosis only in the presence of light (8, 10, 11). It was also shown that ToxA-mediated cell death required ROS production, which was inhibited under dark conditions (41). The observation that *Snn7*-mediated cell death functions in the absence of light indicates that SnTox267 may elicit necrosis from a ROS-independent pathway. Taken together, the identification of SnTox267 has revealed a complex set of host components, some of which may operate cooperatively, involved in the elicitation of PCD.

In the case of necrotrophic effectors, the ability to exploit multiple host targets presents an advantageous scenario for the pathogen. Essentially, it expands the utility of a single necrotrophic effector gene and enables infection on a wider range of host cultivars that may contain one or more of the corresponding susceptibility targets. This evolutionary outcome presents an extreme challenge to wheat breeding programs. Because *SnTox267* is present in the vast majority of North American isolates sampled in this study, the known sensitivity targets *Snn2*, *Snn6*, and *Snn7* need to be prioritized for removal from elite breeding material to effectively combat this disease. Additionally, due to the apparent cooperative nature of *Snn2* and *Snn6*, the possibility also exists that the crossing of two insensitive lines with opposite complements of nonfunctional and functional *Snn2*/*Snn6* copies (i.e. *snn2*/*Snn6* × *Snn2*/*snn6*) may unknowingly result in susceptible progeny. Breeding lines can be rapidly phenotyped with purified SnTox267 infiltrations to prevent this scenario and eliminate susceptibility loci from breeding germplasm. As we have demonstrated that SnTox267 can elicit cell death from two pathways, it is conceivable that additional susceptibility loci may exist. We are currently using purified SnTox267 to evaluate global wheat collections to identify additional targets for SnTox267.

Analysis of *SnTox267* within a natural population of *P. nodorum* revealed the largely conserved presence of the gene. The proportion of isolates harboring a functional copy of *SnTox267* is comparable to levels observed for *SnTox1,* which was found in 95.4% of the same population used in this study, as well as in 84% of isolates in a global collection (20, 21). The prevalence of both *SnTox267* and *SnTox1* are substantially higher than that of *SnToxA* and *SnTox3* which were present in 63.4% and 58.9% of isolates in the collection used in this study, respectively (21). These differences may be attributed to multiple roles of the specific effector protein. For example, the presence of SnToxA within a *P. nodorum* population was found to be highly correlated with the presence of the host sensitivity gene *Tsn1* in local wheat germplasm (21). The strong selection pressure exerted by the SnToxA-Tsn1 interaction likely drives the maintenance of *SnToxA* within the population, however, a fitness penalty may exist as it is readily removed in the absence of *Tsn1*. Another factor contributing to its absence is likely the lack of an identifiable beneficial alternate function. In contrast to the levels of PAV observed for *SnToxA*, the presence of *SnTox267* in nearly all isolates examined suggests it may possess an advantageous secondary function. Although it remains to be experimentally validated, a chitin-binding or other beneficial effector function would explain the propensity to maintain this gene within the pathogen population at similar levels of prevalence observed with *SnTox1,* which has been shown to bind chitin in the fungal cell wall and protect *P. nodorum* from wheat chitinases (42). Another possible explanation for the conserved presence of SnTox267 is the existence of multiple host targets. Currently, three genetically distinct and polymorphic susceptibility loci functioning in two separate pathways have been identified in wheat germplasm, which increases the probability of the introduction of at least a single pathway capable of inducing SnTox267-mediated necrosis. Also, if germplasm used in breeding programs harbored combinations of *Snn2*, *Snn6*, and *Snn7*, there is a lower probability that all targets are removed throughout the breeding process compared to a single gene. Furthermore, the presence of multiple host sensitivity genes suggests that a gene family, or minimally a structural motif, acts as a common target for SnTox267 and additional sensitivity loci may exist in wheat germplasm. This would further broaden the utility of SnTox267, and therefore its maintenance in the natural population.

Genetic compensation or transcriptional adaptation are phenomena observed in other eukaryotic organisms, including zebrafish and Arabidopsis, although the exact mechanism that controls the compensatory response has not been elucidated (43). The deletion of *SnTox267* led to the serendipitous discovery of altered expression profiles of physically unlinked effector genes, indicating that genetic compensation or transcriptional adaptation may be active in *P. nodorum*. Previous research demonstrated similar compensatory effects following effector disruption in *P. nodorum*. Faris et al. provided the initial evidence that effector gene expression can directly impact the contribution of a given effector – susceptibility gene interaction to disease (44). *SnToxA* was found to be expressed significantly higher in isolate Sn5 compared to isolate Sn4. Genetic dissection of the host interactions with these two isolates revealed the *SnToxA – Tsn1* interaction contributed more to disease following Sn5 inoculations compared to Sn4 inoculations. These results directly link elevated *SnToxA* expression to a more substantial impact on disease via interaction with *Tsn1*. Deletion of an effector gene has also been shown to modify unrelated effector expression and subsequently alter host interaction outcomes. Inoculation of a RIL population segregating for host susceptibility genes *Tsn1*, *Snn1*, and *Snn3* with *P. nodorum* isolate Sn2000 (produces SnToxA and SnTox1) resulted in the detection of three QTL, including the expected *Tsn1* and *Snn1* loci. However, inoculation with a Sn2000 *SnToxA*-disrupted mutant led to the detection of *Snn1* at a greater magnitude, explaining substantially more phenotypic variation than inoculations with the wild-type isolate. It was subsequently discovered that *SnTox1* was expressed at a significantly higher level in the knockout isolate compared to the wild-type, indicating that upregulation of *SnTox1* led to a greater contribution of *Snn1* to disease in this population (45). Similarly, a wheat mapping population derived from a cross of varieties Calingiri and Wyalkatchem was observed to segregate for sensitivity to SnTox1 and SnTox3 using infiltration of purified proteins (46). However, inoculation of this population with Australian isolate SN15, which possesses functional copies of both *SnTox1* and *SnTox3*, detected a QTL corresponding to *Snn1* but failed to identify the QTL corresponding to *Snn3*. However, the *Snn3* QTL was detected upon inoculation with a *SnTox1* disrupted mutant of SN15. Furthermore, it was discovered that expression of *SnTox3* was significantly higher in the mutant than in the wild-type isolate, indicating that the deletion of *SnTox1* relieved a suppression mechanism acting on *SnTox3* (46). In our study, the deletion of *SnTox267* appeared to significantly enhance the expression of *SnToxA*, *SnTox1*, and *SnTox3* through some form of genetic compensation. One possible explanation is that a common transcription factor may be regulating all four of these effector genes. Previously, Rybak et al. (47) demonstrated that the zinc-finger transcription factor *PnPf2* regulated both *SnToxA* and *SnTox3*, but did not appear to regulate *SnTox1* expression. If a transcription factor activates *SnToxA*, *SnTox1*, *SnTox3*, and *SnTox267* expression and acts in a dose-dependent manner, the loss of a binding site via deletion of the *SnTox267* genomic region could result in increased binding to promoters of the three remaining effectors and subsequent higher expression. Alternatively, a *trans*-acting repressor, such as a small RNA molecule, may be encoded at this genomic locus and when deleted, lifted suppression of the other effector genes.

These results demonstrate that SnTox267 is a proteinaceous effector that exploits two host pathways to elicit cell death and cause disease. This effector is largely conserved throughout a natural population of *P. nodorum,* suggesting the presence of an advantageous secondary effector function or the presence of multiple host targets beyond Snn2, Snn6, and Snn7. Deletion of *SnTox267* resulted in the compensatory upregulation of three unlinked effector genes, adding more evidence to the existence of a complex regulatory mechanism.

## Materials and Methods

### *Variant identification in a* P. nodorum *natural population*

A natural population consisting of 197 *P. nodorum* isolates was used in this study and was previously described and sequenced by Richards *et al.* (2019). Isolates were collected from wheat producing regions in the United States, including 105 collected from the Upper Midwest, 84 collected from the Southeastern United States, and 8 from the Pacific Northwest. Genome sequencing, as well as single nucleotide polymorphism (SNP) and insertion/deletion (InDel) identification was conducted in a previous study (21) and the derived genotypic data was used in the present research. Briefly, the genome of each isolate was sequenced to approximately 28x coverage. Sequencing reads for each isolate were trimmed using trimmomatic (48) and mapped to the Sn4 reference genome (24) using BWA-MEM (49). SAMtools ‘mpileup’ was used to identify variants (50). Genotypic data for the natural population was retrieved from https://github.com/jkzrich/pnodorum_popgen.

### Phenotyping of differential wheat lines

Inoculum was produced from each *P. nodorum* isolate as previously described (51). Briefly, isolates were cultured on V8 potato dextrose agar (150 mL V8 juice, 3 g CaCO_3_, 10 g Difco PDA, 10 g agar, 850 mL H_2_O). Dried agarose plugs (stored at −20 °C) were streaked across the plate and incubated at room temperature under constant fluorescent light. After approximately seven days, pycnidia formation was observed and spores were harvested by flooding the plate with sterile H_2_O followed by agitation with a sterile inoculation loop. Spores were quantified with a hemocytometer and the concentration was adjusted to 1 × 10^6^ spores/mL. The spore suspension was amended with two drops of Tween20 per 100 mL of suspension.

Each *P. nodorum* isolate was inoculated onto differential wheat lines BG223 (harbors the *Snn2* susceptibility gene) and ITMI37 (harbors the *Snn6* susceptibility gene). Three seeds of each wheat line were planted in cones and represented a single replicate. The moderately susceptible wheat cultivar Alsen was planted as a border to reduce edge effect. Plants were grown under greenhouse conditions until they reached the second leaf stage (approximately two weeks) and inoculated as previously described (51). Briefly, inoculum was applied evenly to the plants using a paint sprayer until runoff was observed. Inoculated plants were then moved to a mist chamber at 100% humidity for 24 hours. Following incubation, plants were moved to a growth chamber set at 21 °C with a 12-hour photoperiod. Disease was scored seven days post-inoculation using a 0-5 rating scale where 0 is highly resistant and 5 is highly susceptible (52). At least three replicates were conducted for each individual *P. nodorum* isolate.

### Association mapping and candidate gene identification

A genome-wide association analysis was conducted using TASSEL (53). Polymorphic sites with greater than 30% missing data or a minor allele frequency less than 5% were removed. A general linear model using the first three components of a PCA as a fixed effect to correct for population structure was tested. P-values were adjusted using a Bonferroni multiple comparison correction and loci with an adjusted p-value of less than 0.001 were considered significant. Candidate genes within 5 kb regions flanking significant markers were identified from the Sn4 genome annotation (https://github.com/jkzrich/pnodorum_popgen). Secretion signals in candidate genes were predicted with SignalP 5.0 (54) and EffectorP (25) was used to predict effector proteins. The DiANNA 1.1 web server was used to predict disulfide bond formation between cysteine residues (55)

### Development of gene-disrupted mutants

The *SnTox267* gene was disrupted using a split-marker approach as described previously (56). Primers SnTox2_KO_F1 and SnTox2_KO_R1 were designed to amplify a 981 bp region upstream of the start codon and primers SnTox2_KO_F2 and SnTox2_KO_R2 were designed to amplify a 1010 bp region downstream of the stop codon (Supplementary Table 3). A 24-nucleotide sequence complementary to overlapping hygromycin resistance gene cassette HY and YG fragments was added to the 5’ ends of the SnTox2_KO_R1 and SnTox2_KO_F2 primers, respectively, to facilitate overlap PCR. PCR reactions were made to amplify the upstream and downstream regions using the following parameters: initial denaturation at 95 °C for 5 min, 95 °C for 30 s, 62 °C for 30 s, and 72 °C for 60 s for a total of 35 cycles, and a final extension at 72 °C for 7 min. The hygromycin resistance gene cassette fragments were amplified from the pDAN plasmid (8) using primer combinations M13F/HY to produce the HY fragment and YG/M13R to produce the YG fragment. The same cycling parameters were used as described above with the exception of an extension time of 2 min. PCR products were purified using the GeneJet PCR Purification kit (ThermoFisher Scientific). A total of 2 μL each of the purified upstream genomic region and HY amplicons were combined with 48 μL of a PCR master mix without terminal primers. A fusion reaction was conducted to facilitate annealing and extension of the overlapping fragments with the following parameters: initial denaturation at 95 °C for 4 min, 95 °C for 30 s, 43 °C for one min, and 72 °C for three min, for a total of four cycles. Following the fusion reaction, 1 μL of each SnTox2_KO_F1 and HY primer (10 μM) were added to the reaction and thermocycling conducted as follows: initial denaturation at 95 °C for 5 min, 95 °C for 30 s, 62 °C for 30 s, and 72 °C for 3 min for a total of 40 cycles, and a final extension at 72 °C for 7 min. The same procedure was followed to produce the fusion product of the downstream genomic region and the YG fragment using the primers YG and SnTox2_KO_R2. Fusion products were purified as previously described and used for fungal transformation. Fungal protoplasting and transformation was conducted as previously described (11).

### Generation of gain-of-function transformants

The *SnTox267* genomic region was cloned into the pFPL-Rh vector as described previously with minor modifications (57). Briefly, the primers SnTox2_DONR_F and SnTox2_DONR_RS were designed to amplify the *SnTox267* locus including 1288 bp upstream containing the putative promoter and terminating with the endogenous stop codon. The primers contained full length attB sequences on the 5’ ends to facilitate Gateway cloning into the pDONR vector. The 2086 bp was amplified from Sn4 genomic DNA in the following reaction: 2 μL template (~50 ng), 0.25 μL Q5 polymerase (New England Biolabs), 5 μL Q5 buffer, 5 μL GC enhancer, 0.5 μL dNTPs (100 μM), 1.5 μL SnTox2_DONR_F1, 1.5 μL SnTox2_DONR_RS, and 9.25 μL H_2_O. PCR parameters were used as follows: initial denaturation at 95 °C for 5 min, 95 °C for 30 s, 62 °C for 30 s, and 72 °C for 2 min for a total of 35 cycles, and a final extension at 72 °C for 7 min. The PCR product was purified as previously described. The streamlined procedure of Gateway cloning previously described by Gong et al. (57) was used to clone the *SnTox267* genomic region into pFPL-Rh. The resulting construct was linearized with *PmeI* and approximately 5 μg of linearized plasmid was used for transformation of *P. nodorum* isolate Sn79-1087. Fungal protoplasting and transformation was conducted as previously described (11).

### Phenotyping of host populations

Two recombinant inbred line (RIL) populations were phenotyped to determine if deletion of *SnTox267* resulted in the loss of interaction with a corresponding host susceptibility gene. The BG population segregated for *Snn2* and the ITMI population segregated for *Snn6*. Inoculum production and inoculation procedures for the gene disrupted mutants and Sn79-1087 gain-of-function transformants was conducted as previously described. For inoculations of the gene disrupted mutants on both populations, two independent mutants were evaluated. Additionally, the Sn4 wild-type and an ectopic mutant containing an intact *SnTox267* and the hygromycin resistance gene inserted elsewhere in the genome were used as controls. Three replications of each population were conducted. For inoculations of the gain-of-function transformants, the Sn79-1087 wild-type and a positive transformant were used. Two replications of each population were conducted.

### Development of a Chinese Spring x CS-Timstein-2D F_2_ population

An F_2_ population derived from a cross between Chinese Spring (CS) and CS-Tm 2D was used for the study. Chinese Spring is a common wheat landrace. CS-Tm 2D is a disomic chromosome substitution line in which the CS chromosome 2D is substituted by the hard red spring wheat cultivar Timstein (Tm) chromosome 2D (16). CS-Tm 2D is sensitive to SnTox267 whereas CS is insensitive. A total of 94 F_2_ plants were infiltrated with SnTox2 at the second leaf stage as previously described (9). The fully expanded secondary leaves of the plants were infiltrated using a 1 ml syringe with the needle removed and the boundaries of the infiltration sites were marked before water-soaking disappeared. Leaves were scored as insensitive or sensitive based on the presence or absence of necrosis within the infiltrated area 5 days after infiltration. A score ≤ 1.0 was considered as an insensitive phenotype and a score > 1.0 was considered as a sensitive phenotype.

DNA was extracted as described by Faris et al. (58). Primer pairs that are polymorphic between CS and CS-Tm 2D were selected from the previously published wheat chromosome 2D map (16). Four markers known to be linked to the *Snn2* gene (59) and five markers known to be linked to the *Snn7* gene (16) were used for genotyping. The PCR conditions were initial denaturation at 94 °C for 5 min, followed by 35 cycles of 94 °C for 30 s, the annealing temperature of 61 °C for 30 s with a 0.2 °C decrement at every cycle, and extension at 72 °C for 90 s, followed by a final extension of 72 °C for 7 min. Amplicons were electrophoresed on 6 % polyacrylamide gels, stained with GelRed stain, and visualized with a Typhoon 9500 variable mode imager (GE Healthcare Life Sciences, Piscataway, NJ). The Kosambi mapping function was used for the linkage analysis using the computer program MapDisto v.1.8 (60). Composite interval mapping was used for QTL mapping using the computer program Qgene 4.0 (61).

### Infiltration of Sn79+SnTox267 culture filtrates in leaves of host populations

Culture filtrates of the Sn79+SnTox267 transformant were used to infiltrate leaves of host mapping populations to map sensitivity to secreted SnTox267. Cultured filtrates of the Sn79+SnTox267 transformant were prepared as previously described (9). Individuals from the BG and ITMI RIL populations, as well as a CS × CS-Tm-2D F_2_ population, were grown in the greenhouse until the second leaf stage. Culture filtrates were infiltrated into the secondary leaves of each line and then placed in a climate-controlled growth chamber. Infiltrated leaves were observed until three days post-infiltration and scored as presence or absence of necrosis.

### QTL analysis of the BG and ITMI populations

Interval mapping (IM) was conducted using the R/qtl ‘scanone’ function with the Haley-Knott regression method. A LOD threshold at the significance of α = 0.05 was determined through permutation analysis with 1000 iterations for each phenotype separately. All significant QTL identified from CIM were fitted into a multiple QTL model to determine the percentage of variation explained by each locus using the ‘fitqtl’ function (62).

### Population genetic and haplotype analysis

Raw sequencing files were obtained from the NCBI SRA (BioProject PRJNA398070) and used for variant confirmation in the *SnTox267* coding region. Sequencing reads were trimmed and mapped to the Sn4 reference genome as previously described (21). Duplicates were marked using ‘MarkDuplicates’ in Picard (http://broadinstitute.github.io/picard/). The Genome Analysis Toolkit Haplotype caller (63) was used to identify SNPs and InDels within the *SnTox267* coding region for each sample individually specifying ‘—intervals Chr14:67371-68168 – emitRefConfidence GVCF’. Genotypes were determined using ‘GenotypeGVCFs’ with default settings. *De novo* genome assembly of each isolate was completed using SPADES with default settings and automatic k-mer selection (64). BLAST databases were generated for each assembly and subsequently searched for the *SnTox267* coding sequence. Total genomic variants, without filtering for missing data or minor allele frequency, were used to create consensus sequences of the *SnTox267* coding region using SAMtools and BCFtools ‘consensus’ (50). Sequences of the coding region were aligned using CLUSTAL Omega (65) and imported into DNASP v6 for population genetic analysis (66). Nucleotide diversity was calculated in 20 bp sliding windows in 5 bp steps across the coding region.

### Differential effector expression

A time course experiment was designed to test the level of *SnToxA*, *SnTox1*, and *SnTox3* expression in both the wild-type Sn4 and Sn4ΔTox267 backgrounds. Plants of wheat cultivar ‘Grandin’ at the secondary leaf stage (typically 14-day old plants) were inoculated side by side with *P. nodorum* isolates Sn4 and Sn4ΔTox267#5. Three samples of leaf tissue for each treatment were collected at five timepoints including 24 h, 48h, 72h, 96h, and 120h post inoculation. Total RNA was extracted using the RNeasy plant mini kit (Qiagen) following the manufacturer’s instructions. RNA was quantified using a Qubit fluorometer. From each sample, 300 ng of RNA was used to synthesize cDNA using the GoScript™ Reverse Transcription System (Promega). Gene specific primers for *Actin* (Pn_Actin_qPCR_F and Pn_Actin_qPCR_R), *SnToxA* (SnToxA_QPCR_F and SnToxA_QPCR_R), *SnTox3* (8981qPCRF and 8981qPCRR) and *SnTox1* (SnTox1_qPCR_F and SnTox1_qPCR_R) were used to perform qPCR for each sample with two technical replicates and three biological replicates (8, 10, 11). Effector gene expression was normalized against expression of the housekeeping gene *Actin*. The entire experiment was conducted three times. Significant differences in gene expression were determined using a t-test (p<0.05).

## Acknowledgements

The authors would like to thank Danielle Holmes for technical assistance. Funding for the project was provided by NIFA-AFRI Competitive Grant no. 2016-67013-24813 from the USDA National Institute of Food and Agriculture.

