## Supplementary figures and images for "A triple threat: the *Parastagonospora nodorum* SnTox267 effector exploits three distinct host genetic factors to cause disease in wheat"

### Supplemental Figure 1

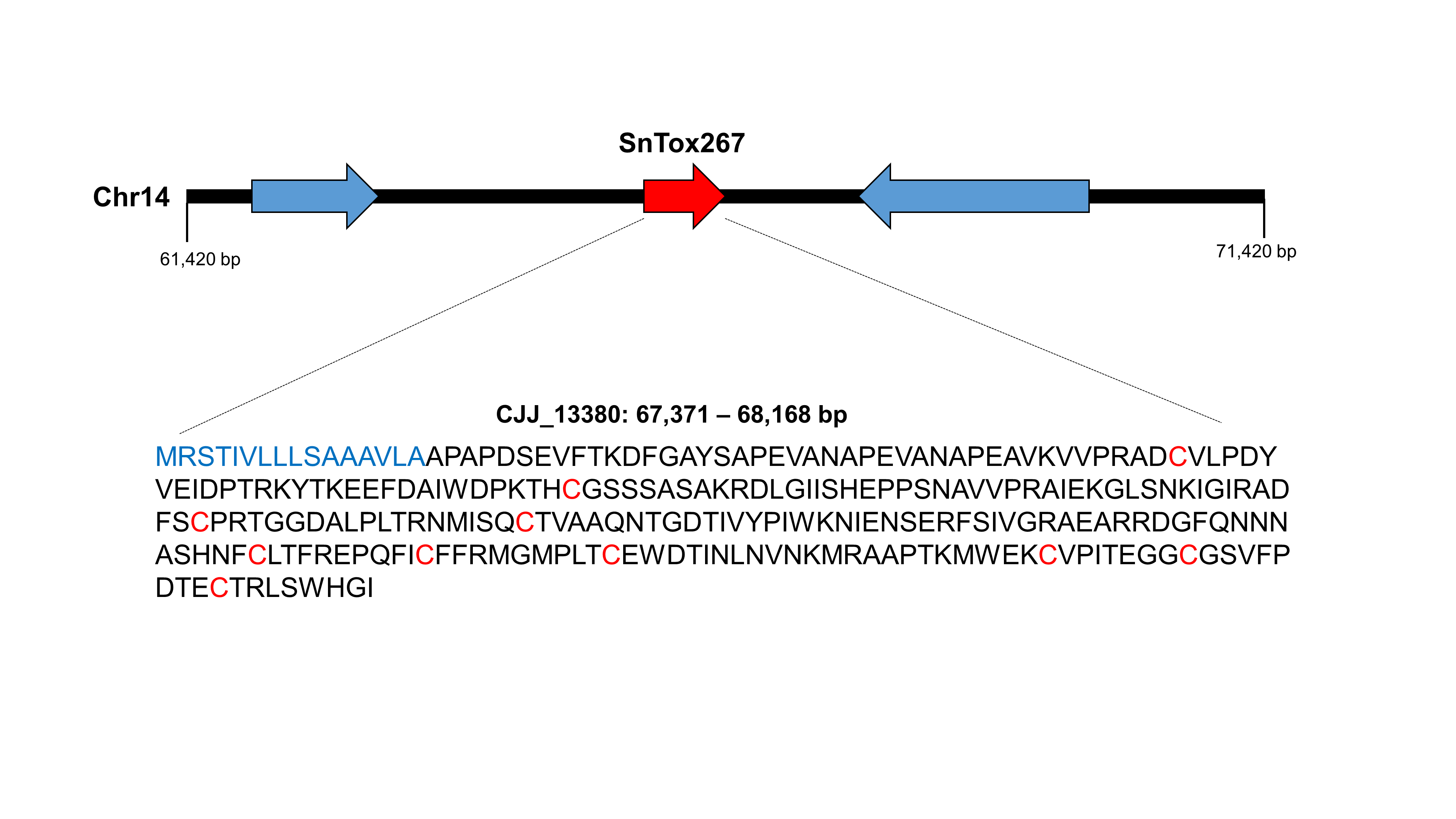

### Supplemental Figure 2

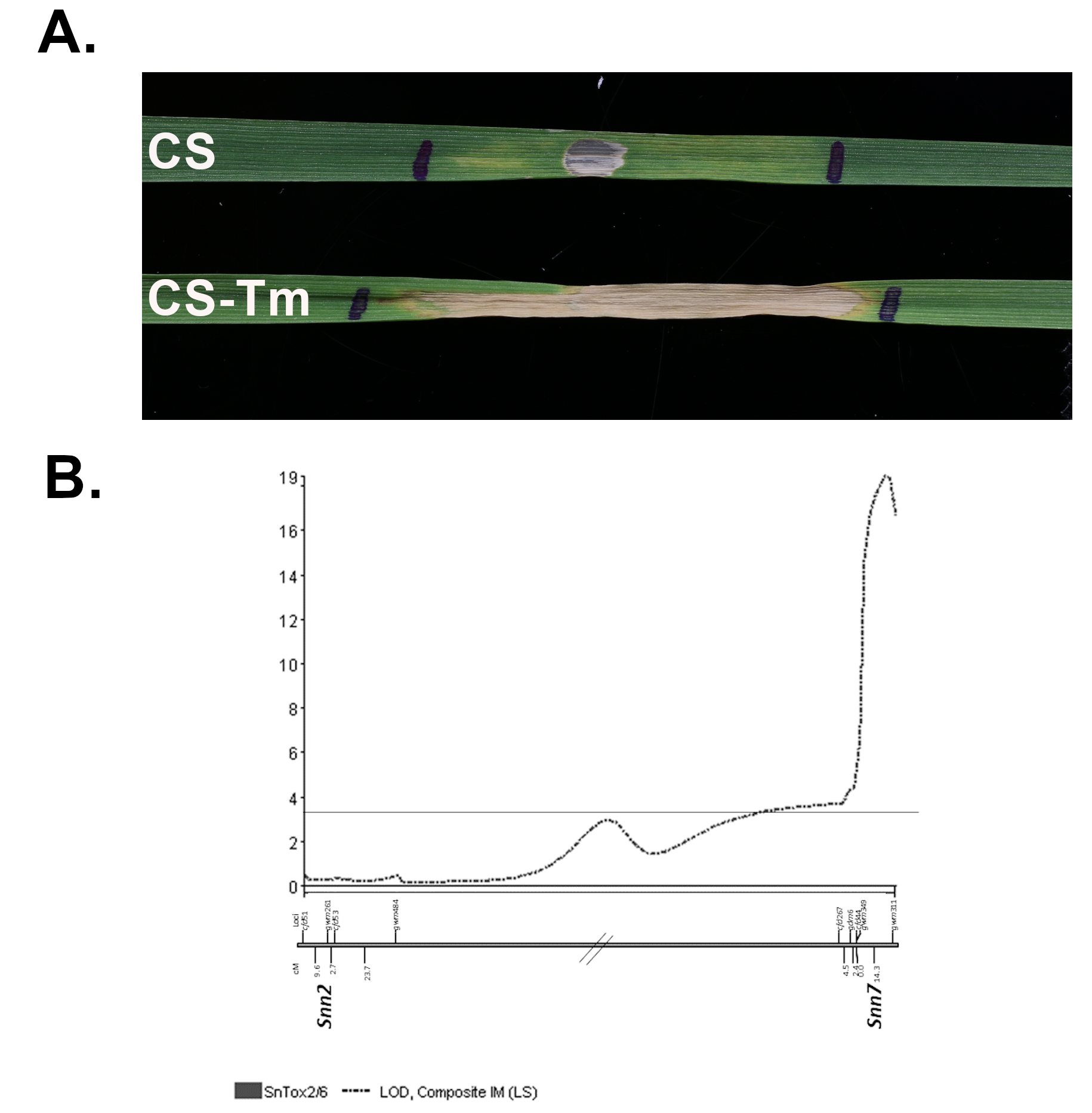

### Supplemental Figure 3

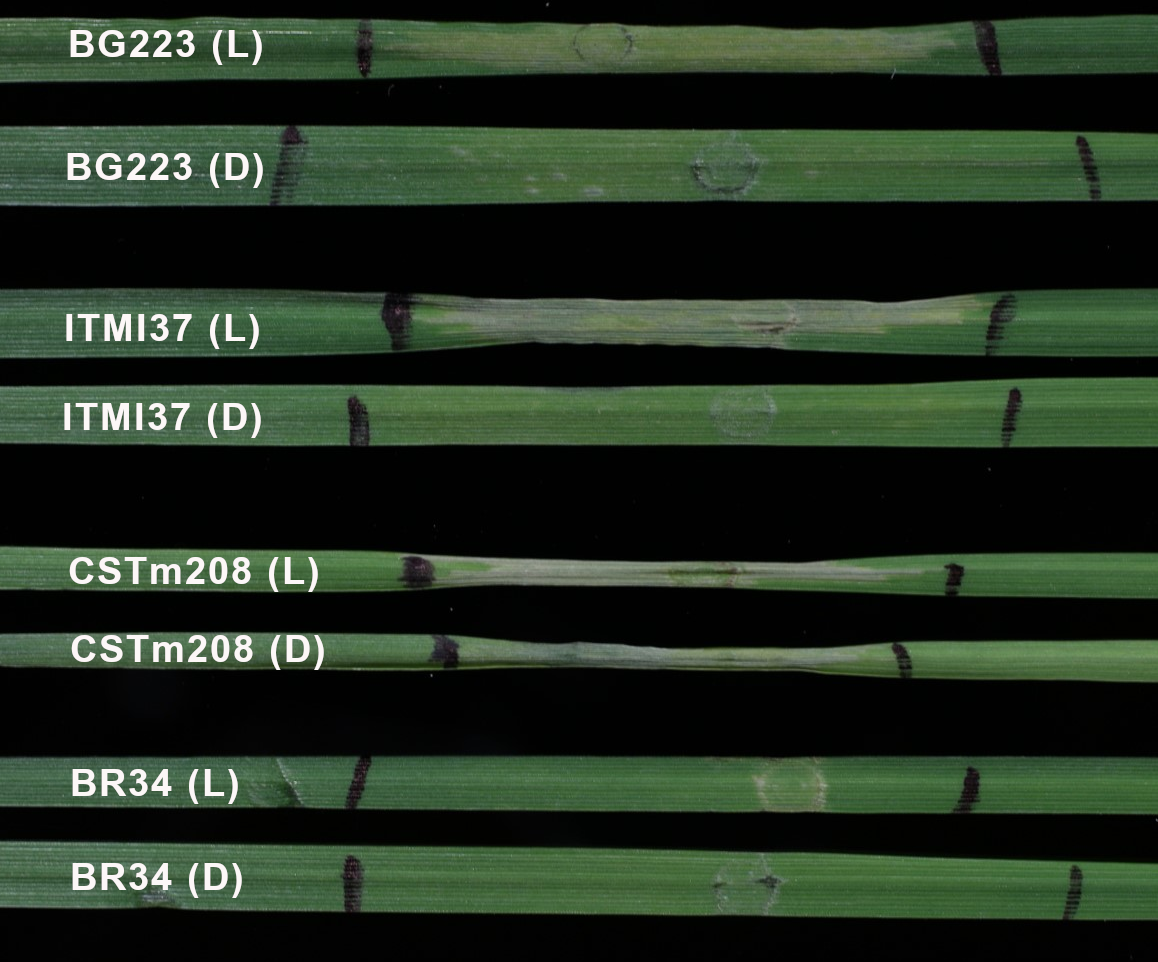

### Supplemental Figure 4

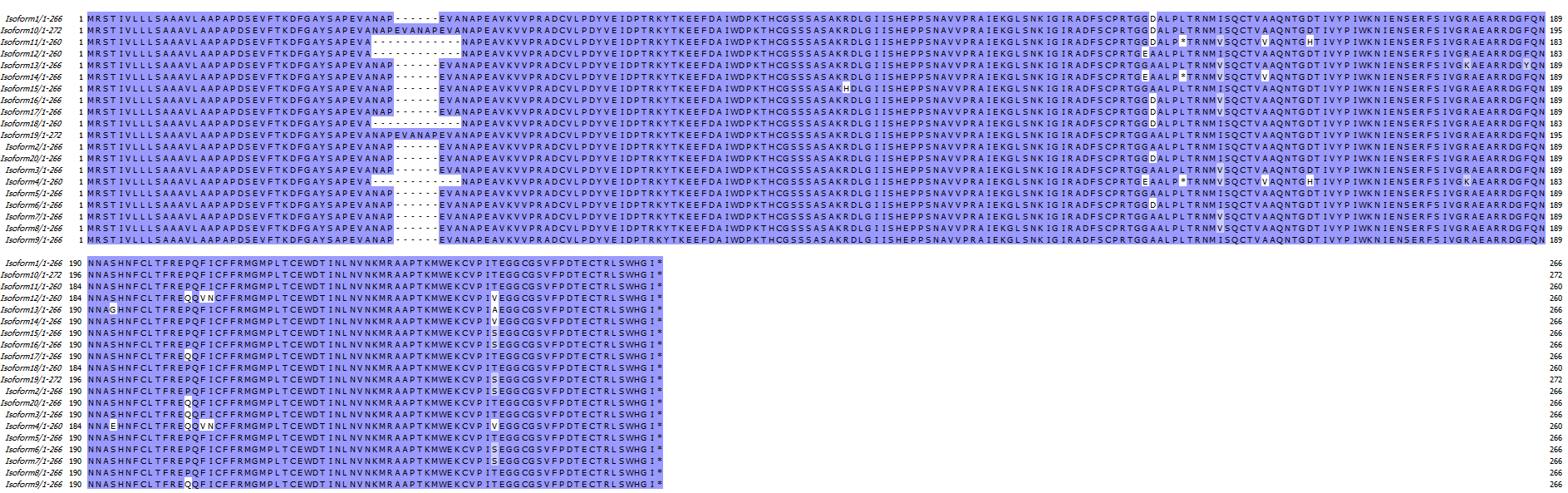

### Supplemental Figure 5

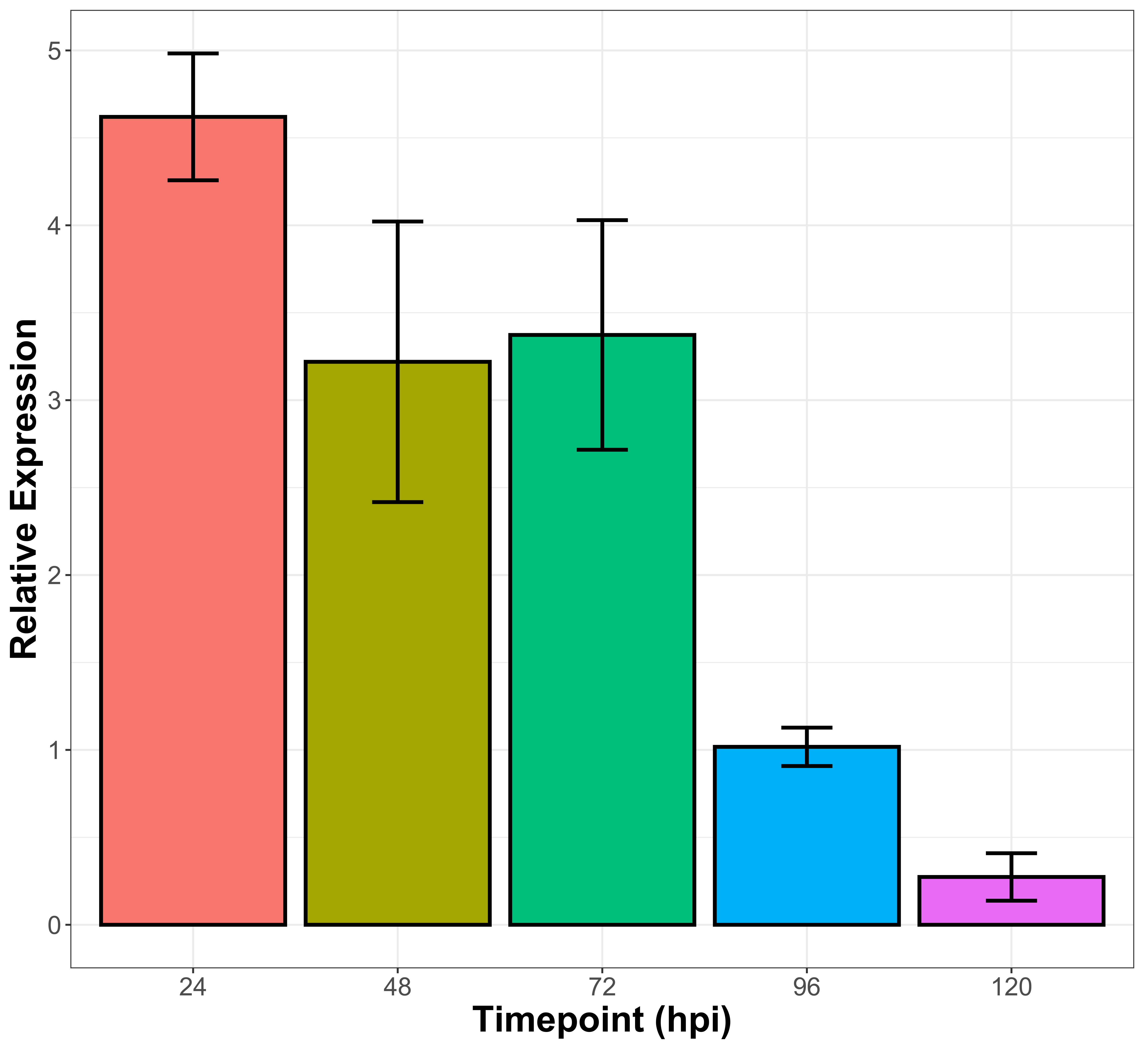
