## Supplemental Figure legends for "A triple threat: the *Parastagonospora nodorum* SnTox267 effector exploits three distinct host genetic factors to cause disease in wheat"

**Supplementary Figure 1.** A 10 kb genomic locus on chromosome 14 underlying the significant association of SNP_66420 with virulence on BG223 and ITMI37. (Top) Solid arrows represent genes within this region and direction indicates orientation. The red arrow represents the *SnTox267* gene, encoding a predicted effector protein. (Bottom) Amino acid sequence of SnTox267. Amino acid residues in blue font represent the predicted secretion signal. Cysteine residues are represented by a red font.

**Supplementary Figure 2.** SnTox267 targets *Snn7* to elicit necrosis. **A.** Infiltration of Sn79_CJJ13380 culture filtrates on wheat lines Chinese Spring (CS) and Chinese Spring – Timstein 2D substitution (CS-Tm) line. **B.** QTL analysis of a Chinese Spring x Chinese Spring – Timstein 2D substitution F_2_ population. The genetic map of wheat chromosome 2D is shown on the x-axis and the logarithm of the odds (LOD) scores are shown on the y-axis. The horizontal line represents a LOD threshold of p=0.05 based on 1000 permutations.

**Supplementary Figure 3.** Snn2 and Snn6 mediated cell death are induced by SnTox267 and are light dependent, whereas Snn7 mediated cell death induced by SnTox267 is light independent. Wheat lines BG223 (*Snn2*), ITMI37 (*Snn6*), CSTm208 (*Snn7*), and BR34 (insensitive parent of the BG population) were infiltrated with culture filtrates of isolate Sn79+13380. ‘L’ signifies that infiltrated plants were kept under normal light conditions following infiltration, whereas ‘D’ signifies that plants were kept under constant dark conditions.

**Supplementary Figure 4*.*** Amino acid alignment of 20 SnTox267 isoforms. Dark shaded boxes represent conserved residue positions. Lightly shaded and white boxes represent amino acid substitutions. ‘-‘ represents a deletion.

**Supplementary Figure 5**. qPCR experiment indicates that *SnTox267* expression peaks at 24 hours post-inoculation (hpi) and declines until 120 hpi. The timepoint (hpi) is shown on the x-axis and the expression values relative to the housekeeping actin gene are shown on the y-axis. Error bars represent the standard error of the means.
